# Phosphatidylserine (PS)-targeting chimeric Interferon (IFN) fusion proteins for anti-tumor applications

**DOI:** 10.1101/2025.01.24.634764

**Authors:** Varsha Gadiyar, Viralkumar Davra, Rachael Pulica, Trevor Frederick, Christopher Varsanyi, Ahmed Aquib, Ziren Wang, Sergey Smirnov, Samhita Bapat, David Calianese, Alok Choudhary, Sergei V Kotenko, Raymond B. Birge

## Abstract

In viable healthy cells, membrane phospholipids are asymmetrically distributed across the lipid bilayer, whereby the anionic phospholipid phosphatidylserine is virtually all distributed on the inner leaflet of the plasma membrane. During apoptosis, phospholipid asymmetry collapses and PS is externalized to the external leaflet where it serves as an “eat-me” signal for efferocytosis, the process whereby dying cells are engulfed and degraded by phagocytes. PS is also externalized on viable activated tumor endothelial cells, stromal cells and cancer cells in the tumor microenvironment reflecting a pathophysiological state of solid cancers that function to suppress host anti-tumor immunity. Several strategies have been envisioned to target dysregulated PS in the tumor microenvironment including PS binding proteins such as Annexin V and PS-targeting monoclonal antibodies (Bavituximab) with promising preclinical results. Here, in an attempt to enhance the efficacy of PS-targeting therapeutics, we have generated a series of recombinant chimeric fusion proteins that fuse type I and type III IFNs (IFN-β-IFN-λ) into a single polypeptide chain separated by a short linker. The IFN-β-IFN-λ fusion proteins retain functions of both type I and type III IFNs but show combined effects to improve biological function as well as enhance anti-tumor activities. To localize IFNs to sites of externalized PS, we next fused the IFN-β-IFN-λ chimeric protein to the PS-targeting gamma-carboxyglutamic acid-rich (Gla) domain of Growth Arrest Specific factor 6 (Gas-6), rendering these IFN biologics as PS targeting modalities. Gas6-IFN-β-IFN-λ proteins selectively bind PS as evident by solid-phase ELISA assays as well as bind PS-positive cells, including apoptotic cells and cells that express CDC50 subunit mutant of the ATP11C flippase. *In vivo*, Gas6-IFN-β-IFN-λ retain strong anti-tumor activities in a syngeneic model when expressed ectopically in a E0771 breast cancer model and B16-F10 melanoma models. Collectively, we report on the generation and utility of a series of novel in class IFN fusion proteins that target the immune stimulatory features of IFNs to the PS externalization in the tumor microenvironment.

Graphical abstract
Gas6-IFN-β-IFN-λ (VitK) have tri-functional activities, acting on a diverse set of cell types to induce an anti-tumor affect. The Gas6 domain aids in homing to the PS rich tumor microenvironment, binding to apoptotic or live stressed PS positive tumor cells. The Gla domain directly binds to PS, whereas the EGF domains help in oligomerization and signal amplification resulting from intermolecular disulphide bonds. The IFN-β domain acts on immune cells such as dendritic cells and macrophages, inducing an interferon response, whereas the IFN-λ domain acts on the tumor epithelial cells, inducing tumor intrinsic anti-tumor activity.

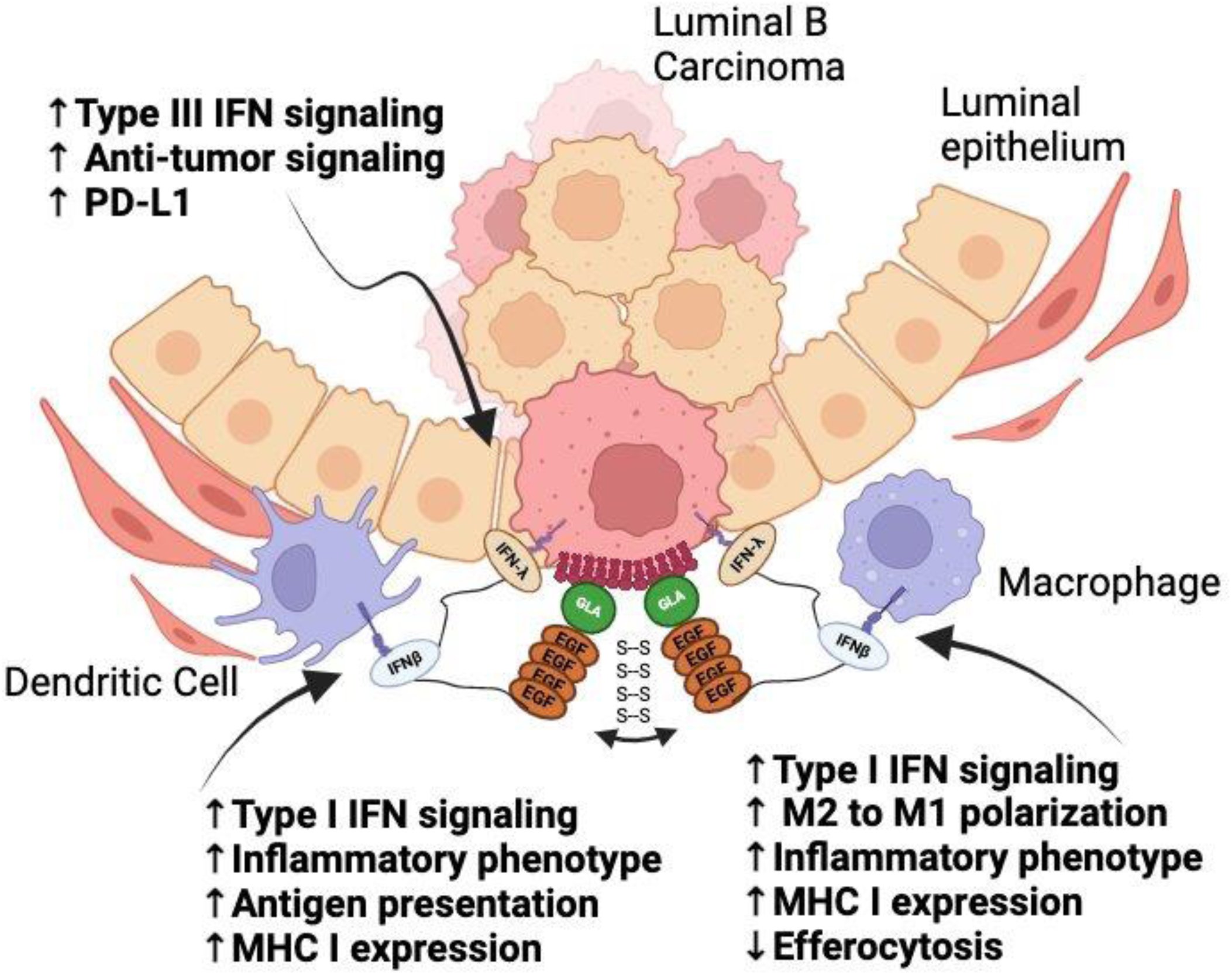

## Introduction

Despite exciting advances in cancer immunology and immuno-therapy, particularly in the conceptualization of immune checkpoint inhibitors such that target inhibitory receptors such as PD1 (Pembrolizumab/Keytruda and Nivolumab/Opdivo) and CTLA-4 (Ipilimumab/Yervoy and Tremelimumab/Imjuno), there is still a great need to identify new treatment options that target generalized epitopes in solid cancers^1^ ^2^ ^3^. Such therapeutic pitfalls and limitations of the checkpoint inhibitors and immune modulators is due in part to the diverse genetics in the etiologies of cancer, clonal evolution via changing mutational landscapes, tumor heterogeneity, and drug resistance^4^ ^5^. Moreover, the microenvironments of many solid cancers also adapt and create an immune tolerant milieu that prevents host anti-tumor immunity benefits of checkpoint therapeutics^6^ ^7^ ^8^ ^9^. In addition, the side effects and off target autoimmunity^10^ further necessitate the identification and characterization of new targets and modalities that can stimulate tumor immunity and benefit cancer patients.

Phosphatidylserine (PS) is an anionic phospholipid that is typically restricted to the inner leaflet of the plasma membrane^11^. However, in many cancer cells and stromal cells in the tumor microenvironment, PS is chronically externalized and can be used as a marker for the tumor mass as well as a target for cancer therapy^12^ ^13^ ^14^ ^15^ ^16^. Moreover, unlike protein targets and the glycocalyx which is highly unstable due to the unique mutational landscapes and distinct N and O-linked glycosylation patterns, externalized PS and its re-localization to the outer membrane represents a stable epitope. Plasma membrane asymmetry is primarily maintained by two types of lipid transporters that include flippases and scramblases^17^. Flippases, such as the aminophospholipid translocases ATP11C and ATP11A, mediate ATP-dependent vectorial translocation of PS from the outer membrane to the cytosolic leaflet, while scramblases, such as Xkr8, and TMEM16F facilitate ATP-independent random shuffling of the phospholipids in activated cells^17^. In recent years, the molecular mechanisms for PS externalization on apoptotic cells and during cell activation have become better understood. For example, during apoptosis, caspase-mediated activation of PS scramblase Xkr8 and caspase-mediated inactivation of the flippases ATP11A and ATP11C results in constitutive PS externalization^18^ ^19^ ^20^. On viable cells, calcium and ATP dependent activation of TMEM16F and P2X7 receptors leads to reversible PS externalization that can return to baseline after cell activation^21^ ^22^. However, on diseased and stressed cells, such as in cancer and during virus infection, PS is externalized patho-physiologically by chronic activation of scramblases, that has been linked to host immune escape^12^ ^23^ ^24^. The rationale that continuous heightened exposed PS can drive immune escape is supported by recent studies show that Xkr8 knockout (which prevents PS externalization, or mutation of the TMEM30A which leads to the so-called PS-out tumors can suppress host immune responses and drive tumor progression ^12^ ^25^.

Based on these observations, many efforts have been envisioned to target externalized on tumors, most emblematically recombinant Annexins and PS-targeting antibodies such as Bavituximab with solid pre-clinical success in murine models ^26^ ^27^. More recent efforts to target PS in tumors has also visioned PS-targeting peptoids ^28^, antibody drug conjugates ^29^, FC-Syt-drug conjugate^30^, and SAPC-DOPS^31^. While clearly meritorious, most of these strategies aim to neutralize PS, which is present at high density on the cell membrane. Moreover, most of the current PS targeting approaches do not implement an immunogenic component, thereby limiting their ability to stimulate host immunity. In the present study, we introduce a conceptually new first in class type of PS-targeting modality, by fusing the PS-targeting domain of Gas6 (a GLA-EGF fusion) to a type I and type III IFN duet.

The rationale to use IFNs is based stems from the long-known feature that IFNs in *vivo* represents one the prime components of host natural defense in higher metazoans for viral infections and cancer^32^ ^33^. IFNs are divided into three families, type I, type II and type III, based on sequence homology as well their receptor usage and activities^34^. In humans, type II IFN includes a single member, IFN-γ, which is more divergent than type I and type III IFNs in terms of IFN structures, expression patterns, and biological actions^35^. By contrast, type I and type III IFNs are more functionally comparable; they are induced by similar pathogen sensing pathways, activate analogous post-receptor signaling and transcriptional pathways, and regulate the expression of the identical set of IFN-stimulated genes (ISGs), the IFN signature^36^ ^37^. At the biochemical level, all type I IFNs bind to a common heterodimeric IFNAR receptor, comprised of IFNAR1 and IFNAR2 subunits, while type III IFNs interact with a common heterodimeric IFNLR, comprised of IFNLR1 (also known as IL28RA) and IL10R2 (also known as IL10RB) ^38^ ^39^ ^40^. While IFNAR and IFNLR share structural and biochemical elements enabling the activation of common JAK/STAT and ISGF3 signaling and transcriptional activators by the two distinct ligand-receptor systems ^41^ ^42^ in recent years it has been shown that type I and type III IFNs target *in vivo* differential sets of cell types and tissues. In this respect, while type I IFN receptors are more ubiquitously expressed in a variety of nucleated cells, type III IFN receptors are predominantly expressed in cells lining epithelial barriers that include the respiratory, gastrointestinal, and reproductive tracts ^43^ ^44^ ^45^. These observations indicate functional compartmentalization of the action of type I and type III IFNs and suggest that type III IFNs represent a more restricted first line of defense at sites of viral exposure as well as adenocarcinomas, while type I IFNs regulate a more robust and systemic response.

In addition to their role in antiviral immunity, both type I and type III IFNs appear to have significant anti-tumor activities ^46^ ^47^. Type I IFNs, such as IFN-α and IFN-β, can have direct effects on tumor cells by suppressing proliferation or by inducing apoptosis or necroptosis ^48^ ^49^. Type I IFNs can also act as adjuvant therapeutics to enhance anti-tumor immunity, including the polarization of macrophages towards M1 phenotype, the activation, maturation, and enhanced cross presentation by DCs and the subsequent immune responses towards tumor-associated antigens, as well as direct activation of T and NK cells ^50^ ^51^ ^52^. These features are the antithesis of what occurs in PS-positive solid tumors. Likewise, type III IFNs also appear to have anti-tumor properties and can inhibit proliferation and stimulate apoptosis of tumor cells ^53^ ^54^. Although recent studies indicate that neutrophils and DC subsets are directly responsive to type III IFNs, most immune cells do not express IFNLR1 and are not sensitive to IFN-λs. Therefore, type III IFNs might be better adapted for the use as anticancer biologics as adjuvants to other immunotherapeutics, or in combination with type I IFNs. Recent studies indeed suggest that IFN-α and IFN-λ can have concerted activities to suppress certain cancers such as HCC *in vivo* ^55^. A role for type III IFNs in cancer biology is further supported by findings that IFNLR knockout mice are more susceptible to sarcoma formation induced by carcinogen exposure^56^. Additionally, IFN-λ induces CXCL10 when added to mammary epithelial cells leading to the increased recruitment of T cells to the tumor site ^57^. Collectively, both type I and type III IFNs have both direct and indirect effects on tumor cells and within the tumor microenvironment.

Based on pleotropic activities induced by type I and type III IFNs in distinct subsets of target cells in vivo, we have engineered and generated a series of recombinant IFN fusion proteins that combine type I (IFN-β) and type III (IFN-λ) IFNs into a single polypeptide chain (IFN-β-IFN-λ) to be used as broad-spectrum antiviral agents and in cancer biology. By combining IFN-β-IFN-λ into a single polypeptide with a linker, such chimeric IFNs are expected to act focally to activate immune pathways on epithelial-immune interfaces, thereby enhancing immune outcomes. Additionally, we also fused the IFN-β-IFN-λ with the phosphatidylserine (PS) binding Gla domain of Growth Arrest Specific Factor 6 (Gas6). Gas6 and IFN-β-IFN-λ fusion protein (referred to as Gas6-IFN-β-IFN-λ throughout the paper) are developed to localize fusion proteins to regions of constitutive PS externalization. Here we present a proof-of-concept study describing a first-in-class PS-targeting modality that targets a type I and type III IFN modality with immune-oncology applications.

## Material and Methods

### Cell culture

Mouse triple negative breast cancer E0771 cells (CH3 Biosystems LLC), mouse melanoma B16-F10 cells, human non-small cell lung carcinoma H1299 cells, mouse lung adenoma LA4 cells, human retinal epithelium ARPE19 cells, and mouse intestine epithelium cells (mIECs) were maintained in RPMI-1640 medium (Sigma-Aldrich), HEK293T cells in DMEM medium and CHO IFN-λR reporter cells were cultured in HAM’s F12 medium. All media were supplemented with 10% fetal bovine serum (FBS; Atlanta biologics), 100 IU/ml penicillin and 100 μg/ml streptomycin (Sigma-Aldrich). Cells were grown at 37°C in a humidified incubator containing 5% CO_2_. After thawing, cells were used for up to 8-10 passages and their authenticities were checked by Short Tandem Repeat (STR) analysis according to manufacturer’s protocol (GenePrint 10 System, Promega). Cells were routinely checked for mycoplasma contamination.

### Expression plasmids for IFN fusion proteins

Novel IFN fusion proteins were cloned into expression vectors pEF2 vector (EF-1α promoter) or pcDNA3.1 vector (CMV promoter) using infusion cloning. The fusion proteins were designed to contain a flexible linker (ASGSSGGSSGTSGSSGGSSGTST) between the IFN-β and IFN-λ, and between the Gas6(G+E) and IFN-β domains. The proteins also express a 6X His tag at the c-terminal for purification.

#### IFN fusion protein production

HEK293T cells were transfected with PEF2 vector encoding various IFN or Gas6-IFN fusion expression constructs (7 μg plasmid per 10 cm plate at ∼80% confluency using 28 μl LipoD293 transfection reagent). 24 hour post-transfection, the cells were washed 3 times with medium without serum and replaced with either medium alone or medium containing either 2 μg/ml vitamin K (Phytonadione injectable emulsion from Hospira) or 2 μM Warfarin (Sigma). Cell culture supernatants containing secreted IFN proteins were collected at 72 h post-transfection. For mock, cells were transfected with empty vector and cell supernatant collected as above. Secretion of fusion molecules was confirmed by anti-His immunoblotting as all the proteins are His-tagged.

Purified recombinant IFN fusion proteins were produced by transfecting Expi293T cells (ThermoFisher, Cat: A25869) with corresponding expression vectors. 18 h post-transfection, cells were supplemented with 2 μg/ml vitamin K, cell culture supernatants were collected at 96 h post-transfection, and the proteins were purified via TALON resin. 1.0 μg purified protein was subjected to SDS-PAGE under denaturing conditions and purity was assessed by Coomasie Blue staining.

#### Immunoblotting

Whole cell lysates were prepared in HNTG buffer (20 mM HEPES, pH 7.5, 150 mM NaCl, 10% glycerol) supplemented with 1% Triton X-100, 1 mM PMSF, 1 mM sodium orthovanadate, 10 mM sodium molybdate, 1mM EDTA, 10 mM sodium fluoride and 1% protease inhibitor cocktail. Cells were lysed, scraped, incubated on ice for 10 min and centrifuged at 12,000 X rpm for 10 min. Cleared lysates were collected, mixed with SDS containing Laemmli buffer, boiled for 5 min and resolved by SDS-PAGE. Immunoblotting was performed with monoclonal antibody (mAb) against either β-actin (mouse mAb; Cell Signaling, MAB374), or phospho-STAT1 (rabbit mAb; BD Bioscience, 612233) as primary Abs followed by secondary horseradish peroxidase (HRP)-conjugated Affinipure Goat anti-mouse Ab (Jackson ImmunoResearch, 115-035-166) or anti-rabbit Ab (Jackson ImmunoResearch, 115-035-144), respectively.

#### Antiviral Assay

Cells were seeded in a 96 well plate and treated with serially diluted (a one-third dilution) amounts of either recombinant murine IFN-α or IFN-λ2 or IFN-containing cell culture supernatants starting with 100 ng of recombinant IFN proteins or 50 ml of cell supernatants in the first well in duplicates. After 18 h of IFN treatment, vesicular stomatitis virus (VSV) was added and incubated for 24 or 48 h, after which cells were fixed, and cell viability was determined by staining cells with 0.1% crystal violet solution followed by reading absorbance at 590 nm.

#### PS ELISA

96-well ELISA plates (Greiner bio-one) were coated with L-PS (brain, porcine, Avanti Polar Lipids, Cat: 840032P and 840012) dissolved in methanol (12.5 μg/ml; 100 ml/well) by placing the the plates in a sterile fume hood at 22°C until methanol evaporated. ELISA plates were then blocked with 5% BSA in PBS overnight at 4°C. PS-coated plates were washed 3 times with lipid-wash buffer (LWB: 10 mM HEPES, 150 mM NaCl, 2.5 mM CaCl_2_) and incubated with soluble proteins (IFN-β-IFN-λ, γ-carboxylated or non-γ-carboxylated Gas6-IFN-β-IFN-λ) diluted in 5% BSA and titrated 2-fold (the concentration range: 10 to 0.16 μg/ml) for 1 h at 22°C. Plates were then washed 3 times with LWB and PS-bound proteins were detected by ELISA performed with His mAb (ThermoFisher, Cat: MA1-21315; 1:1000 dillution; 1 h at 37°C) followed by alkaline phosphatase (AP)-conjugated anti-mouse IgG (Jackson ImmunoResearch, Cat: 115-055-146; 1:2000 dilution; 1 h at 37°C) and addition of diethanolamine (DEA) buffer containing AP substrate (Sigma, Cat: S0942; 1 mg/ml). Plates were washed 3 times in PBS to remove unbound primary and secondary Ab and amounts of PS-bound proteins were assessed by measuring absorbance at 405 nm over the course of 2 h.

### Protection assay

96-well ELISA plates (Greiner bio-one) were coated with L-PS (brain, porcine, Avanti Polar Lipids, Cat: 840032P and 840012) dissolved in methanol (12.5 μg/ml; 100 ml/well) by placing the the plates in a sterile fume hood at 22°C until methanol evaporated. ELISA plates were then blocked with 3% BSA in Annexin V Binding Buffer for 1 hour at 37°C. PS-coated plates were washed 3 times with lipid-wash buffer (LWB: 10 mM HEPES, 150 mM NaCl, 2.5 mM CaCl_2_) and incubated with HEK293T supernatants containing IFN-β-IFN-λ, γ-carboxylated or non-γ-carboxylated Gas6-IFN-β-IFN-λ diluted in 3% BSA and titrated 3-fold for 1 h at 37°C. Plates were then washed 3 times with LWB. mIEC cells were plated on these plates. 48 hours later, after mIEC cells had adhered and formed a monolayer, VSV was added incubated for 48 hours more. Cell viability was determined by staining cells with 0.1% crystal violet solution followed by reading absorbance at 590 nm.

#### Mouse experiments

Immunocompetent female or male C57BL/6 (6-8 weeks old) were obtained from Jackson laboratory. C57BL/6 *Ifnar1^-/-^*, *Ifnlr1^-/-^*, *Ifnar1^-/-^Ifnlr1^-/-^* mice and SCID mice were bred as homozygotes at the facility. All mice were maintained in specific pathogen-free (SPF) barrier facility, maintained under a strict 12 h light-dark cycle with access to regular chow diet and autoclaved reverse osmosis water. For mammary tumor cell inoculation, 0.5 X 10^5^ E0771 cells either mock-transfected (control cells transfected with empty plasmid), or expressing either IFN-β, IFN-λ, IFN-β-IFN-λ, Gas6(Gla)-IFN-β-IFN-λ, Gas6(Gla+EGF)-IFN-β, Gas6(Gla+EGF)-IFN-λ, or Gas6(Gla+EGF)-IFN-β-IFN-λ, or 50:50 combination of E0771 cells expressing IFN-β and IFN-λ, or Gas6(Gla+EGF)-IFN-β and Gas6(Gla+EGF)-IFN-λ were re-suspended in RPMI-1640 complete medium containing Matrigel (50 % v/v; Corning) and injected into the 9/10 mammary fat pad of female C57BL/6 mice (n=8/group). For melanoma model, 0.1 X 10^6^ B16F10 cells either moc-transfected or expressing IFN-β-IFN-λ and Gas6(Gla+EGF)-IFN-β-IFN-λ were injected subcutaneously into flank of male C57BL/6 mice. The tumor growth was assessed twice a week by caliper measurement of tumor diameter in the longest dimension (*L*) and at right angles to that axis (*W*). Tumor volumes were estimated using the formula *L* X 2*W*.

The total body weight of mice was measured once a week until the end of the study. On day 29, mice were euthanized with CO_2_ inhalation; and primary tumors, lungs and spleens were collected for further analysis. Lungs were washed once in PBS and stained using bouin solution for metastases quantification. Metastasis incidences were calculated by counting metastatic nodules in the lungs under magnification microscope. Mouse experiments were performed in accordance with the guidelines and under the approval from Rutgers Institutional Animal Care and Use Committee at New Jersey Medical School.

### Cell Surface Binding

W3/CDC50^-/-^ cells or calcium ionophore treated H1299 cells (20 mM) were counted and 5 X 10^5^ cells were used for staining. Cells were incubated with either HEK293T cells supernatants of fusion proteins or 20 mg/mL of purified proteins in Annexin V binding buffer at 4C for 15 minutes. After washing, they were stained with His-PE antibody for 15 minutes. The cells were washed 3 times and binding was analyzed by flow cytometry.

### RNA sequencing

MLE-15 cells were cultured in a 6-well plate (1×10^6^ cells/well) and treated (1 ng/ml) with Gas6-IFN-β-λ2 (VitK) or left untreated. RNA was collected at 7 hours using the Qiagen RNeasy Plus Mini Kit according to the manufacturer’s instructions. RNA quantity and purity were assessed using a Nanodrop spectrophotometer. RNA samples were then submitted to the Rutgers Genomics Core for sequencing. RNA-Seq analysis was done in R (4.3.1). Gene set enrichment analysis was performed in R with the Bioconductor package clusterProfiler (version 3.18) where all gene ontologies (Biological Processes, Molecular Functions, Cellular Components) were included in the analysis, and the dot plot was generated with the Bioconductor package DOSE (version 2.10.6). Volcano plots were generated with TidyVerse package ggplot2 (version 3.5.0), where significance was determined by a p-value < 0.05 and log2FC > 1 or log2FC < -1.

#### Statistical analysis

All *in vitro* and *in vivo* experiments were repeated at least three times. Differences between groups in all *in vivo* experiments were examined for statistical significance using a two-tailed Student’s t-test and one-way ANOVA to compare multiple groups. For virus infection experiments, data were analyzed using two-way ANOVA followed by statistical significance analysis using Sidak’s multiple comparisons test for weight loss studies, and one-way ANOVA followed by statistical significance analysis using Tukey’s multiple comparisons test. GraphPad Prism software was used to perform statistical analysis. P<0.05 was considered significant.

## Results

### Cloning and characterization of recombinant type I/type III IFN fusion proteins

While type I and type IIII IFNs both activate the JAK/STAT/ISGF3 signaling pathway and induce similar sets of target genes that activate antiviral and anti-tumor immunity, expression studies indicate that type I receptors (IFNAR) and type III IFN receptors (IFNLR) are functionally compartmentalized^58^. In this capacity, type III IFN receptors are expressed on mucosal barrier-covering epithelial cells (lung, gastrointestinal, reproductive tracts) while type I IFN receptors are more ubiquitously expressed on almost all nucleated cells, the latter representing a more systemic and robust immune activation that the local activation of type III IFN receptors (**Fig.1.a.**)^58^. Here, in an attempt to improve efficacy and targeting of IFNs, we engineered a series of recombinant fusion proteins that fuse type I IFN (IFN-β) and type III IFN (IFN-λ), separated by linker, into a single polypeptide chain (**Fig. 1.b.**) with the goals to focally activate both Type I and Type III IFN responding cells at immune cell/epithelial cell boundaries.

**Figure 1.**
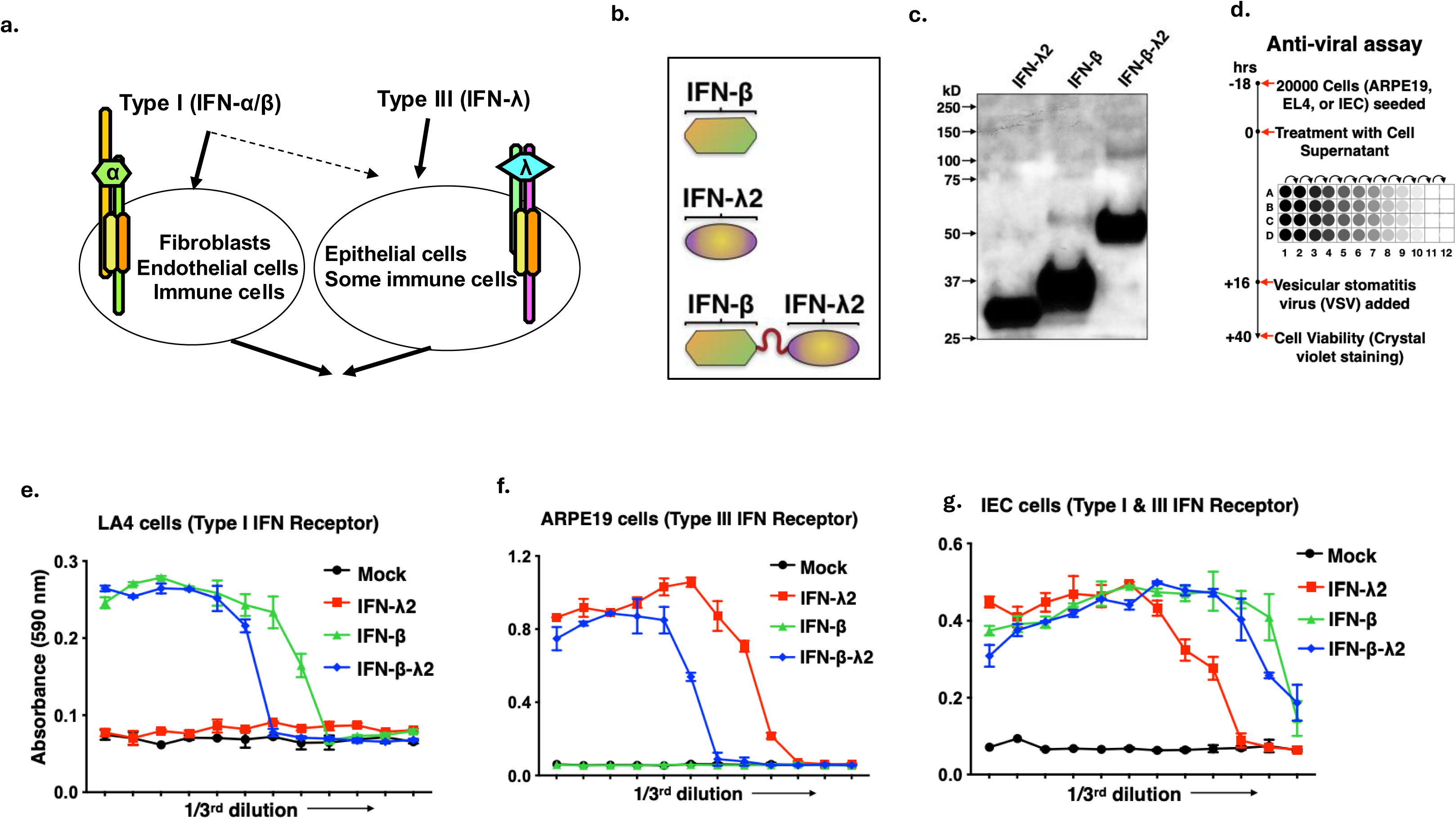
Design, production and characterization of novel IFN fusion proteins. **(a)** Action of IFNs are compartmentalized, where type I IFNS act on fibroblast, endothelial cells and most immune cells, whereas type III IFNs act mainly on epithelial cells. **(b)** The schematic illustration of IFN fusion proteins**. (c)** Immunoblot showing secretion of each protein at their expected molecular weight in the cell supernatant when probed using anti-his antibody. **(d)** The schematic illustration depicting antiviral assay strategy. The IFN receptor expressing cells were pretreated with HEK293T cell supernatant containing IFN, IFN fusion and Gas6-IFN fusion proteins. After 12 h of pretreatment, cells were treated for 24 h with vesicular stomatitis virus (VSV) and cell viability was determined by staining cells with 0.1% crystal violet solution followed by reading absorbance at 590nm**. (e)** The representative graphs showing the antiviral activity of IFNs in LA4, mouse lung adenoma cells (express type I IFN receptor) **(f)**, ARPE19, the human retinal epithelium cells (respond only to mouse type III IFNs) **(g)**, and mIEC, mouse intestine epithelium cells (express both type I and type III IFN receptors).

To first generate type I and type III recombinant proteins for proof-of-concept feasibility studies and to show that chimeric proteins retain functional IFN activities, DNA constructs encoding murine IFN-β, murine IFN-λ, or murine IFN-β-IFN-λ fusion construct were cloned into a PEF2 expression vector with a c-terminal His tag and following transfection into HEK293 cells, recombinant proteins in the culture supernatant were assayed by immunoblotting with anti-His mAb. All proteins were produced at expected molecular weights without notable degradation (**Fig. 1.b, c.**). Subsequently, to test the *in vitro* biological activity of the proteins, we performed antiviral protection assays as illustrated in **Fig. 1.d.** For antiviral assays, we utilized 3 different cell lines depending on the expression of IFN receptors. These included mouse lung adenoma LA4 cells (that express type I IFN receptor) (**Fig. 1.e.**), human retinal epithelium ARPE19 cells (that express to type I and type III IFN receptors) (**Fig. 1.f.)**, and mouse intestine epithelium cells (mIECs) (that express both type I and type III IFN receptors) (**Fig. 1.g.**). To demonstrate antiviral activity, above-mentioned cells were treated with serial dilutions of IFN-containing HEK293 cell culture supernatants (**Fig. 1.d.**). After 18 h of IFN treatment, vesicular stomatitis virus (VSV) was added and incubated for 48 h, after which cells were fixed, and cell viability was determined by staining cells with crystal violet solution followed by reading absorbance at 590 nm. Th e absorbance (cell viability) in the representative graphs depicts antiviral protection by IFN molecules. As expected, the LA4 cells responded only to IFN-β and IFN-β-IFN-λ (**Fig. 1.e.**) while the ARPE19 cells only responded to IFN-λ, and IFN-β-IFN-λ since mouse IFN-λ but not mouse IFN-β can cross react with human IFNLR (**Fig. 1.f.**). The mIECs responded to IFN-β, IFN-λ and IFN-β-IFN-λ proteins **(Fig. 1.g.)**. Together, these data affirm that both IFN-β and IFN-λ moieties within the IFN-β-IFN-λ fusion protein maintain their biological activities and that the fusion molecule containing type I and type III IFN duets are stable and biologically active.

Subsequently, to test if IFN-β-IFN-λ fusion protein display enhanced potencies compared to IFN-β and IFN-λ given together, IEC cells were treated with equivalent amounts of IFN-β alone, IFN-λ alone, IFN-β and IFN-λ in 1:1 ratio and IFN-β-IFN-λ fusion protein. 24 hours later, the cells were infected with VSV. 24 hours later, antiviral potency was measured by staining live cells by crystal violet. While, IFN-β and IFN-λ required approximately 130 pg/mL IFNs to show 50% activity, IFN-β-IFN-λ fusion protein showed 50% activity at very low amounts (**Fig. 2.a.**), indicating enhanced antiviral potency when proteins are expressed as fusion proteins separated by a linker.

**Figure 2.**
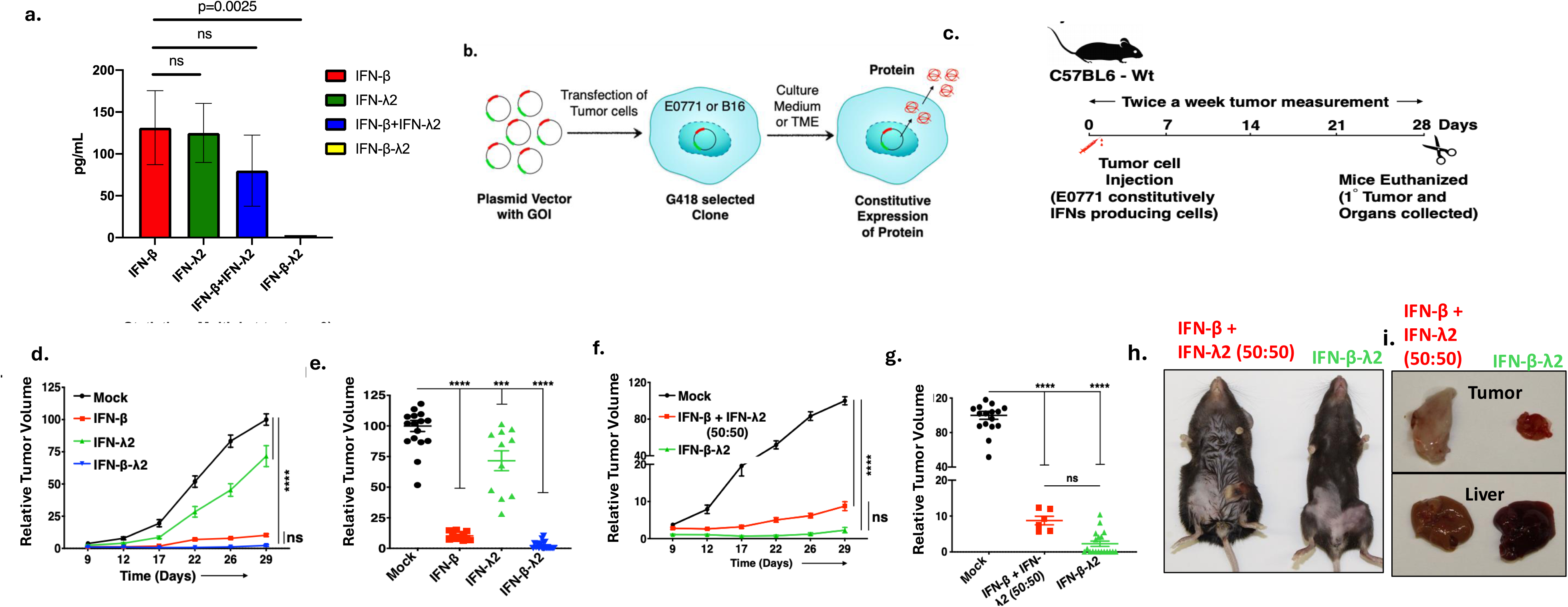
Anti-tumor effect of IFNs and IFN fusion proteins in the E0771 murine breast cancer tumor model. **(a)** Antiviral assay (VSV) at 24 h pre-treatment with equivalent amounts of IFNs on IEC cells. Concentration at which IFN treatment showed 50% activity is plotted on Y axis (p=0.0025, n=3, multiple t-tests). **(b)**The schematic illustration showing transfection of E0771 or B16-F10 cells followed by antibiotic selection to create stable cell lines that constitutively secrete IFN fusion proteins. **(c)** The graphical illustration showing experimental outline of the E0771 murine breast cancer tumor model in C57BL/6 mice. 5 X 10^4^ E0771 cells, mock or constitutively secreting IFN-β, IFN-λ, or IFN-β -IFN-λ, or IFN-β and IFN-λ in 50:50 ratio were injected orthotopically into the mammary fat pad of female C57BL/6 mice. Tumor growth was monitored every 3-5 days for 4 weeks and tumor volumes are shown as relative mean (**d, f**) or individual (**e, g**) tumor volumes compared to the mean volume of mock-transfected tumors measured at the end point (day 29). **(h)** Mice injected with E0771 cells expressing IFN-β and IFN-λ in 50:50 showed ascites, whereas the mice injected with IFN-β -IFN-λ fusion did not show this phenotype.**(i)** Tumors expressing IFN-β -IFN-λ fusion protein, and livers from these mice show reduced anemia compared to the tumors expressing IFN-β and IFN-λ in 50:50.

### Type I and type III IFNs have anti-tumor functions in immunocompetent syngeneic mouse models of tumor growth

To investigate effects of IFN signaling on tumor growth *in vivo*, we used two immunocompetent mouse tumor models, E0771 murine breast cancer model and a B16-F10 murine melanoma model. First, to assess sensitivity of E0771 and B16-F10 cells to type I and type III IFNs, the cells were treated for 18 h with recombinant murine IFN-α and IFN-λ2, infected with VSV and cell viability was measured at 48 h for E0771 cells and 24 h for B16-F10 cells post VSV infection (**Supplementary Figure 1.a, 1.b)**. Antiviral protection was observed in E0771 cells only with IFN-α but not with IFN-λ2, whereas both IFNs have provided antiviral protection in B16-F10 cells. These data indicate that E0771 cells respond only to type I IFNs, whereas B16-F10 cells are sensitive to both type I and type III IFNs, albeit at different capabilities.

Next, to explore the anti-tumor function of IFNs and IFN fusion molecules, we generated stable transfectants of E0771 and B16-F10 cells that constitutively secrete IFNs and IFN fusion molecules (**Fig. 2b**.). No overt toxicity (apoptosis or necroptosis) in the cultured cells was observed nor did cells show decreased cell growth using an MTT assay (**Supplementary Figure 1.d**.). Accordingly, when ectopically IFN secreting modified tumor cells are implanted into syngeneic mice, the tumor cell-produced IFNs act locally in the tumor microenvironment. For the *in vivo* anti-tumor studies, 0.5 X 10^5^ mock-transfected E0771 cells, or E0771 cells constitutively producing IFN-β, IFN-λ, or IFN-β-IFN-λ fusion proteins were orthotopically transplanted into mammary fat pad of C57BL/6 female mice. After tumor cell injection, tumor volume was measured twice a week and body weight once a week over the period of 4 weeks as depicted in the experimental outline **(Fig. 2c.)**. As shown in **Fig 2d-e**, both IFN-β and IFN-β-IFN-λ showed strong anti-tumor activity, while IFN-λ showed a more modest effect. As reported earlier, IFN-β has also shown anti-tumor effect as well, however strong off-target effects such as, loss of hematopoiesis, ascitic fluid accumulation in the peritoneal cavity were observed (data not shown). In agreement with data shown in **Fig. 1.g**., these data indicate that IFN-β-IFN-λ retains functional activity. This is further supported by observations that E0771 tumors grew slower in mice transplanted with the IFN-β-IFN-λ E0771 expressing cells compared to mice transplanted with 50% E0771 IFN-β cells and 50% E0771 IFN-λ cells mixed (**Fig.2.f and g**). Taken together, data in **Fig. 2f-g** support the idea that the combined action of type I and type III IFNs with the IFN fusion molecule has enhanced capacity to modulate the local tumor microenvironment to suppress tumor growth. In addition to marked synergistic anti-tumor activities of IFN-β-λ2, mice with tumors expressing a combination of single IFN molecules developed ascites (**Fig.2.h.)** and anemia, as suggested by pale appearance of the liver and tumor (**Fig.2.i**), whereas mice bearing tumors expressing IFN-β-λ2 lacked these signs, suggesting reduced off target effects of the IFN-β-λ2 fusion molecules.

### Novel Gas6-IFN fusion proteins display both IFN activity and PS binding characteristics

Additionally, to target IFNs to tissues with chronically externalized PS, and transform the tumor from an immuno-suppressive (cold) to a pro-inflammatory (hot) microenvironment, single IFNs and the IFN-β-IFN-λ fusion protein were further fused to the Gla domain or the Gla and EGF-like domains of Gas6 (a PS-binding TAM receptor ligand) creating bi-functional Gas6-IFN fusion proteins (**Fig. 3.a.**). Having shown that recombinant type I and type III IFN fusion molecule can be stably generated and retains the intrinsic biological activities of both IFN moieties, we next fused IFNs, either alone or in tandem, to the Gla domain or Gla plus 4 tandem EGF-like domains of Gas6, hence endowing these proteins to bind externalized PS. We hypothesized that PS binding function of Gas6 will serve as a targeting molecule to bring IFNs in the PS rich tumor microenvironment or virus infected sites. In other iterations of these fusion proteins, phosphorylation labeling sites were introduced to facilitate labelling and in vivo localization (**Supplementary table 1**). PS has been well studied as a global immunosuppressive signal ^[26]^ and targeting PS by utilizing Gas6-IFN fusion molecules is expected to target IFNs to PS positive environment and convert global immunosuppressive signals into IFN-promoted immunostimulatory signals (**Fig.3.a.**). To assess whether Gas6-IFNs also retain PS-binding capabilities, we first prepared fusion proteins by culturing HEK293 cells in the presence of vitamin K or Warfarin, (the latter blocks γ-carboxylation of the Gas6 Gla domain) (**Fig. 3.b.**). as shown by immunoblotting with an anti-Gla antibody.

**Figure 3.**
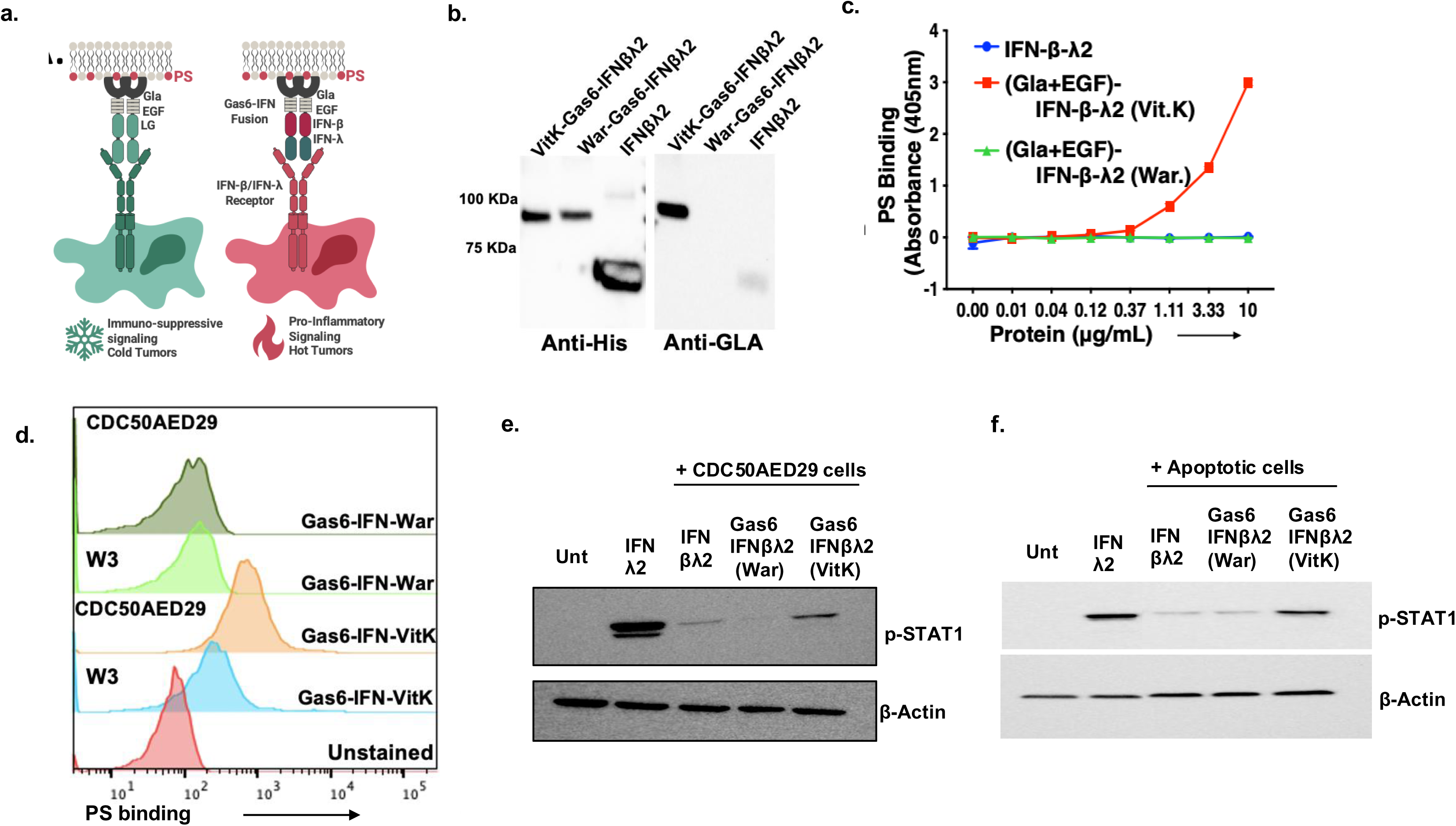
Novel Gas6-IFN fusion proteins display both IFN activity and PS binding characteristics. **(a)** Schematic showing the rationale for developing Gas6(Gla+EGF)-IFN-β-IFN-λ fusion proteins; Gla and EGF domains of Gas6 are required for PS binding, whereas, the LG domain activates immunosuppressive TAM receptor signaling. Truncating the Gas6, and replacing the LG domains with immunomodulatory IFN fusions will convert immunosuppressive signaling to pro-inflammatory signaling. **(b)** γ-carboxylated (PS binding) and non γ -carboxylated (non PS binding) variants of Gas6(Gla+EGF)-IFN-β-IFN-λ were produced by supplementing transfected HEK293T cell media with vitamin K or Warfarin respectively during protein production. **(c)** PS ELISA confirms that γ-carboxylated Gas6(Gla+EGF)-IFN-β-IFN-λ binds to PS whereas IFN-β-IFN-λ does not. **(d)** This is further confirmed by using CDC50AED29 cells, a mouse lymphoma cell line (W3) that harbors a mutation in CDC50, leading to the loss in PS-flippase function and constitutive externalization of PS on live cells. Gas6(Gla+EGF)-IFN-β-IFN-λ (VitK) binds to only CDC50AED29 (PS^+^) cells whereas Gas6(Gla+EGF)-IFN-β-IFN-λ (War) does not bind to either. **(e)** CHO IFN-λ reporter cells were treated with IFN-β-IFN-λ, Gas6(Gla+EGF)-IFN-β-IFN-λ (VitK) and Gas6(Gla+EGF)-IFN-β-IFN-λ (War) with CDC50AED29 cells **(f)** or apoptotic cells for 15 min, washed and centrifuged to remove unbound protein and incubated with reporter cells for 30 min. Immunoblot of pSTAT1 is used as a read out of IFN-λR activation in the reporter cells.

As previous studies have shown that γ-carboxylation is required for PS binding ^[28, 29]^, this biochemical approach permits the development of fusion proteins with and without the ability to bind PS-positive cells, as shown in the solid phase ELISA using affinity-purified recombinant His-tagged proteins (**Fig. 3.c.**). Preferential PS binding was also observed when recombinant proteins were incubated with CDC50AED29 cells, a mutant cell line that fails to internalize PS and that show constitutively externalized PS on live cells (**Fig. 3.d.**) ^[30, 31]^. Similar results were observed when human non-small cell lung carcinoma H1299 cells were treated with calcium ionophore for 15 min to induce PS externalization. Following calcium ionophore treatment, cells were incubated with fusion molecules for 5 min and their binding on the surface of normal and stressed cells was detected with His Ab using flow cytometry. Similar to the PS ELISA, only γ-carboxylated Gas6-IFN-β-IFN-λ was able to bind the externalized PS on the surface of stressed H1299 cells in the pattern similar to Annexin-V **(Supplementary figure 3.f.**) that was used as a positive control for PS binding in flow cytometry. These data demonstrate that under conditions of cell stress, mimicked here by calcium ionophore, Gas6-IFNs bind to cells with externalized PS and therefore are predicted to “home to” a stressed environment with constitutively externalized PS such as a tumor micro-environment. Additionally, in order to assess if replacement of the LG domains of the Gas6 with IFN domains in the Gas6(Gla+EGF)-IFN-β-IFN-λ fusion proteins have an effect on the efferocytosis capability of these modified Gas6 ligands, we treated BMDMs with apoptotic Jurkat cells in presence of the fusion proteins. The IFN fusion proteins were seen to significantly reduce efferocytosis of apoptotic cells by BMDMs, indicating a potential anti-tumorigenic effect (**Supplementary Fig.3.g.)**

To better substantiate this latter idea, we employed a CHO IFN-λ reporter cell line, which expresses a modified IFN-λ receptor complex composed of the intact IL-10R2 and the chimeric IFN-λR1/IFN-γR1 chain where the IFN-λR1 extracellular domain is fused to the intracellular IFN-γR1 domain. Accordingly, in the presence of IFN-λ2, the reporter receptor will dimerize and activate IFN-γR1-specific signaling leading to phosphorylation of STAT1 as a read out (**Fig. 3 e,f**). As indicated by the detection of pSTAT1, these reporter cells respond to fusion proteins containing IFN-λ, including IFN-β-IFN-λ and Gas6-IFN-β-IFN-λ fusion proteins prepared in the presence of vitamin K or Warfarin, when treated with IFNs with or without apoptotic cells for 30 minutes. To reiterate PS binding function, fusion proteins were incubated with CDC50A^ED29^ cells (**Fig.3.e.**) or apoptotic cells (**Fig.3.f**.) for 5 mins followed by centrifugation and cell pellet was washed twice with DMEM medium to remove unbound proteins. Subsequently, cell pellet containing CDC50A^ED29^ cells or apoptotic cells was used to induce STAT1 phosphorylation in the IFN-λ reporter cells. As shown by immunoblot analysis, pSTAT1 was detected only with γ-carboxylated Gas6-IFN-β-IFN-λ protein bound to CDC50A^ED29^ cells (**Fig. 3.e**) or apoptotic cells **(Fig. 3.f.)**. A priori, these data indicate that γ-carboxylated Gas6-IFN fusion proteins can simultaneously bind PS and at the same time activate IFN receptors.

### Gas6-IFN-βλ2 (Vit-K) displays unique clustered pattern of binding on PS^+^ cells

To explore the effects of the addition of the Gas6 domain to the IFNβ-IFN-λ fusion proteins, we developed several iterations of this protein, with either the Gla domain only (Gas6 (Gla)-IFN-β-IFN-λ, Gas6 (Gla)-IFN-β, Gas6 (Gla)-IFN-λ,) or Gas6 containing both the Gla and EGF domains, (Gas6 (Gla+EGF)-IFN-λ, Gas6(Gla+EGF)-IFN-β, and Gas6 (Gla+EGF)-IFN-β-IFN-λ) (**Fig.4.a.).** When these proteins were expressed in HEK293 cells (cultured in the presence of Vitamin K to allow γ-carboxylation of the Gas6 Gla domain), both Gas6(Gla)-IFNs and Gas6 (Gla+EGF)-IFNs were produced, as shown by immuno-blotting with anti-His (**Fig.4.b.)** and effectively γ-carboxylated as evident by immunoblotting with mAb against a γ-carboxylated Gla domain (**Fig.4.c**.). Interestingly, proteins that contained the 4 tandem EGF-like domains appear as both monomers and dimers suggesting that the EGF-like domains control a level of oligomerization (**Figs. 4.b. and 4.c).** This EGF mediated clustering of PS binding Gla proteins can also be observed by amplification of PS receptor (Mertk) signaling, in the presence of a PS source such as apoptotic cells or PS liposomes (**Fig.4.d.).** When Mertk-**γ**R1 reporter cells (Mertk-γR1 reporter construct contains extracellular and transmembrane regions of Mertk and the intracellular kinase domain of IFNγ receptor) are treated with full length Gas6 (Gla+EGF+LG) (**Fig.3.a**.), Mertk is hyperactivated in the presence of PS, provided by apoptotic cells or PS liposomes, perhaps by the clustering of the Gas6 ligands (**Fig.4.d**). Having shown that Gas6-IFN-β-λ2 (VitK) preferentially binds to PS on W3 - CDC50A^ED29^ cells, we explored the pattern of surface binding on these cells. For this, we used imaging flow cytometry or AMNIS. Cells were prepared as they were for flow cytometry, and images collected at 60 X magnification on ImageStream AMNIS. After gating on the singlet cell population, dead cells were gated out using 7AAD staining (**Supplementary figures 3.a-3.c.).** Interestingly, Gas6-IFN-β-λ2 (VitK)-His-PE showed a unique pattern of binding wherein the fluorescence appeared to be clustered or polarized on the cell surface (**Fig.4.e, 4.g**), whereas, a more diffuse binding pattern in the IFN-β-λ2-His-PE samples, (**Fig.4.f, 4.g**), perhaps indicating a pattern of binding to the IFN receptors on these cells. Further analysis showed that the “diffuse staining” population was enriched in the cells treated with IFN -β-λ2-His-PE (**Fig.4.h.)**, whereas the “polarized staining” population was enriched in the cells treated with Gas6-IFN-β-λ2 (VitK)-His-PE (**Fig.4.i.**). Gas6-IFN-β-λ2 (War)-His-PE did not have many positive cells to analyze, since they did not show preferential PS binding (**Supplementary Fig.3.e**.) This unique pattern of polarized staining in the PS dependent binding of Gas6-IFN-β-λ2 (VitK)-His-PE supports the idea that Gla+EGF domains are driving the clustering of the PS molecules, as well as to other Gas6-IFN-β-λ2 (VitK) fusion proteins by engaging certain inter-molecular bonds.

**Figure 4.**
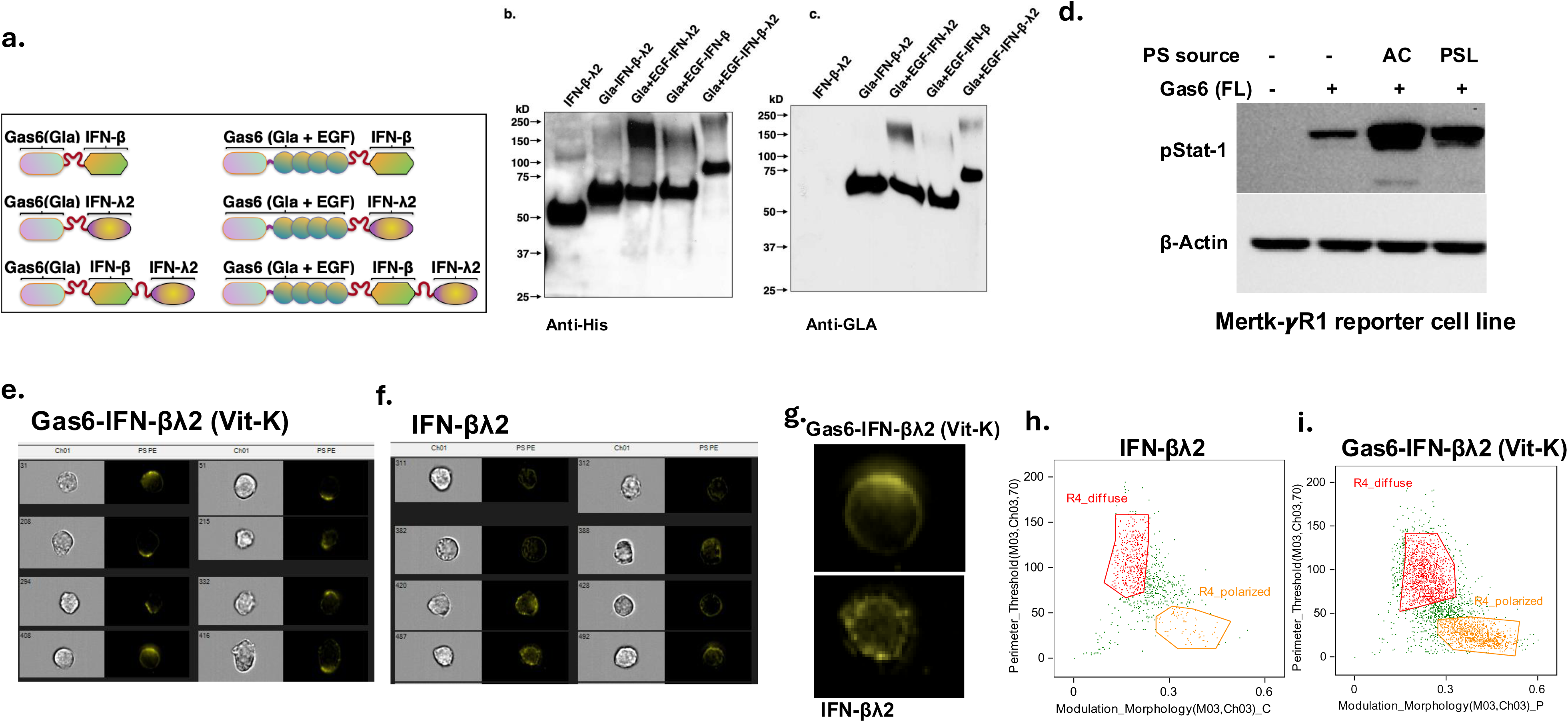
Anti-tumor effect of IFN-β-IFN-λ and Gas6-IFN-β-IFN-λ fusion molecules. **(a)** Schematic diagram showing the different Gas6-IFN fusion proteins developed with just the Gla domain or the Gla and EGF domains of Gas6 with IFN-β, IFN-λ, or IFN-β-IFN-λ i.e. Gas6(Gla)-IFN-β, Gas6(Gla)-IFN-λ, Gas6(Gla)-IFN-β-IFN-λ, and Gas6(Gla+EGF)-IFN-β, Gas6(Gla+EGF)-IFN-λ, Gas6(Gla+EGF)-IFN-β-IFN-λ. **(b)** Immunoblot showing secretion of each protein at their expected molecular weight in the cell supernatant when probed using anti-His antibody and confirming the γ-carboxylation on the Gla domain using anti-Gla antibody **(c). (d)** Activation of Mertk-γ-R1 reporter cell line using Gas6 and PS shows that addition of PS in the form of apoptotic cells or PS liposomes shows hyperactivation of Mertk. Imaging flow cytometry shows distinct pattern of fusion protein binding to CDC50AED29 (PS^+^) cells. **(e)** Gas6-IFN-β-IFN-λ (VitK) shows clustered pattern of binding, whereas **(f)** IFN-β-IFN-λ2 shows a more diffuse pattern of staining. **(g)** shows enlarged images. **(h)** Quantification of the diffuse vs the clustering pattern shows more diffuse pattern with the IFN-β-IFN-λ2 versus, clustered pattern with Gas6-IFN-β-IFN-λ (VitK).

### Anti-tumor effects of Gas6-IFN fusion proteins *in vivo*

In order to test the anti-tumor efficacies in vivo, the E0771 or B16 tumor cells were stably transfected with plasmids containing the above-mentioned constructs, which were then actively secreted by the clonally-derived tumor cells. The E0771 and B16 cells expressing IFN-β-IFN-λ or the Gas6(Gla+EGF)-IFN-β-IFN-λ were themselves protected from virus infection (**Supplementary figure 1.c.).** Analogous to IFN-β and IFN-λ above, all fusion proteins displayed antiviral protection on respective cell lines that sensitive to only mouse type I IFNs (LA4 cells) (**Supplementary figure 2.a.**) or mouse type III IFNs (ARPE19 cells) (**Supplementary figure 2.b**.). This demonstrates that fusion of monomeric or dimeric IFNs to Gas6 Gla or Gla-EGF domains retain their intrinsic biological activities of IFNs as Gas6-IFN fusion chimeric proteins.

Interestingly, while Gas6(Gla)-IFN-β-IFN-λ, Gas6(Gla+EGF)-IFN-λ, Gas6(Gla+EGF)-IFN-β all showed partial reduction in tumor volume, the Gas6(Gla+EGF)-IFN-β-IFN-λ showed a more robust effect and completely blocked tumor growth. This suggests that the IFN-β-IFN-λ duet is the more optimal combination of IFNs, but also that the EGF-like domains of Gas6 also likely participate in signaling possibly by stabilizing the protein or inducing a form of oligomerization (**Fig. 5.a, 5.b.**). This is further supported by the observations shown in **Fig. 5.c, 5.d** that 50:50 mixture of cells expressing either Gas6(Gla+EGF)-IFN-β or Gas6(Gla+EGF)-IFN-λ did not phenocopy the tumor growth profile cells expressing the Gas6(Gla+EGF)-IFN-β-IFN-λ fusion protein. To expand these observations to a second tumor model, we compared IFN-β-IFN-λ and Gas6 (Gla+EGF)-IFN-β-IFN-λ side-by-side in both the E0771 orthotopic model (**Fig. 6.a.,b.**) and the B16-F10 flank model (**Fig. 6.c.,d.**), and showed that Gas6(Gla+EGF)-IFN-β-IFN-λ performed equally in terms of tumor growth suppression as IFN-β-IFN-λ did, again indicating that the fusion of the PS-binding Gla-EGF domains of Gas6 did not interfere with the biological function of the IFN-β-IFN-λ duet as a trimeric protein (**Fig. 6.a. 6.-d.**).

**Figure 5:**
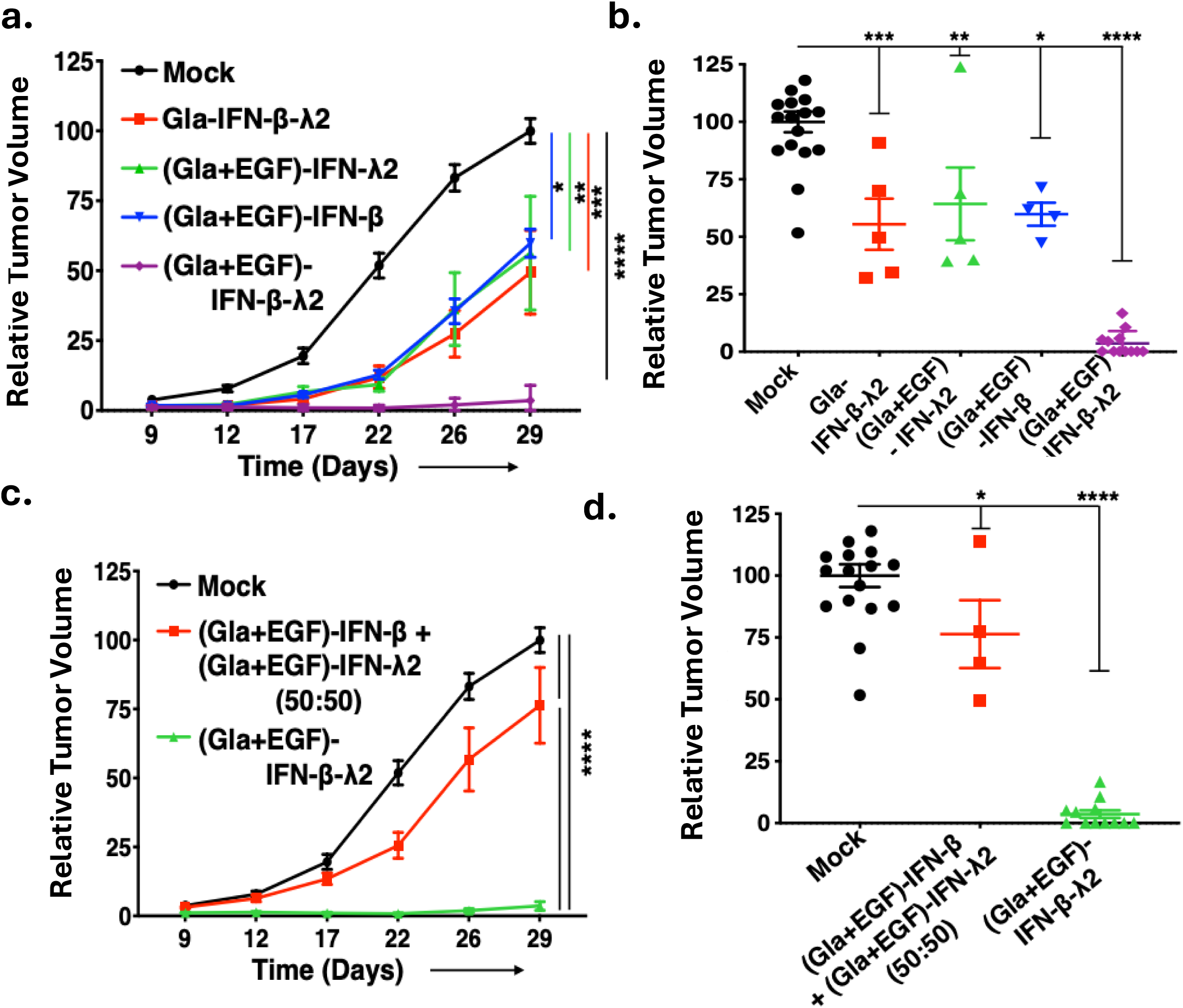
Novel fusion proteins show anti-tumor efficacy in E0771 and B16-F10 tumor models. *In vivo* tumor growth curves of E0771 and B16-F10 tumor models. 5 X 10^4^ E0771 cells, mock transfected, or constitutively secreting either Gas6(Gla)-IFN-β-IFN-λ, Gas6(Gla+EGF)-IFN-λ2, Gas6(Gla+EGF)-IFN-β or Gas6(Gla+EGF)-IFN-β-IFN-λ, were injected orthotopically into the mammary fat pad of C57BL/6 mice and tumor growth determined by tumor volume measurements every 3-5 days for a period of 4 weeks. To compare anti-tumor effects of Gas6-IFN-β-IFN-λ fusion protein to the effects of Gas6-IFN-β and Gas6-IFN-λ (single IFN proteins) given in combination, C57BL/6 mice were implanted with 5 X 10^4^ E0771 cells producing Gas6(Gla+EGF)-IFN-β-IFN-λ cells or a 50:50 mixture of E0771 cells producing Gas6(Gla+EGF)-IFN-β and E0771 cells producing Gas6(Gla+EGF)-IFN-λ (total number of implanted cells 5 X 10^4^). The Gla+EGF domains and the fusion of IFNs β and λ have an additive effect, as shown by reduction in (a) tumor volume by days and (b) tumor volume at end point. Similarly, The Gas6(Gla+EGF)-IFN-β-IFN-λ fusion protein was much more potent in inhibiting tumor growth than the 50:50 combination of single IFNs as shown by relative tumor volume at various time points, and at end point **(c,d).**

**Figure 6:**
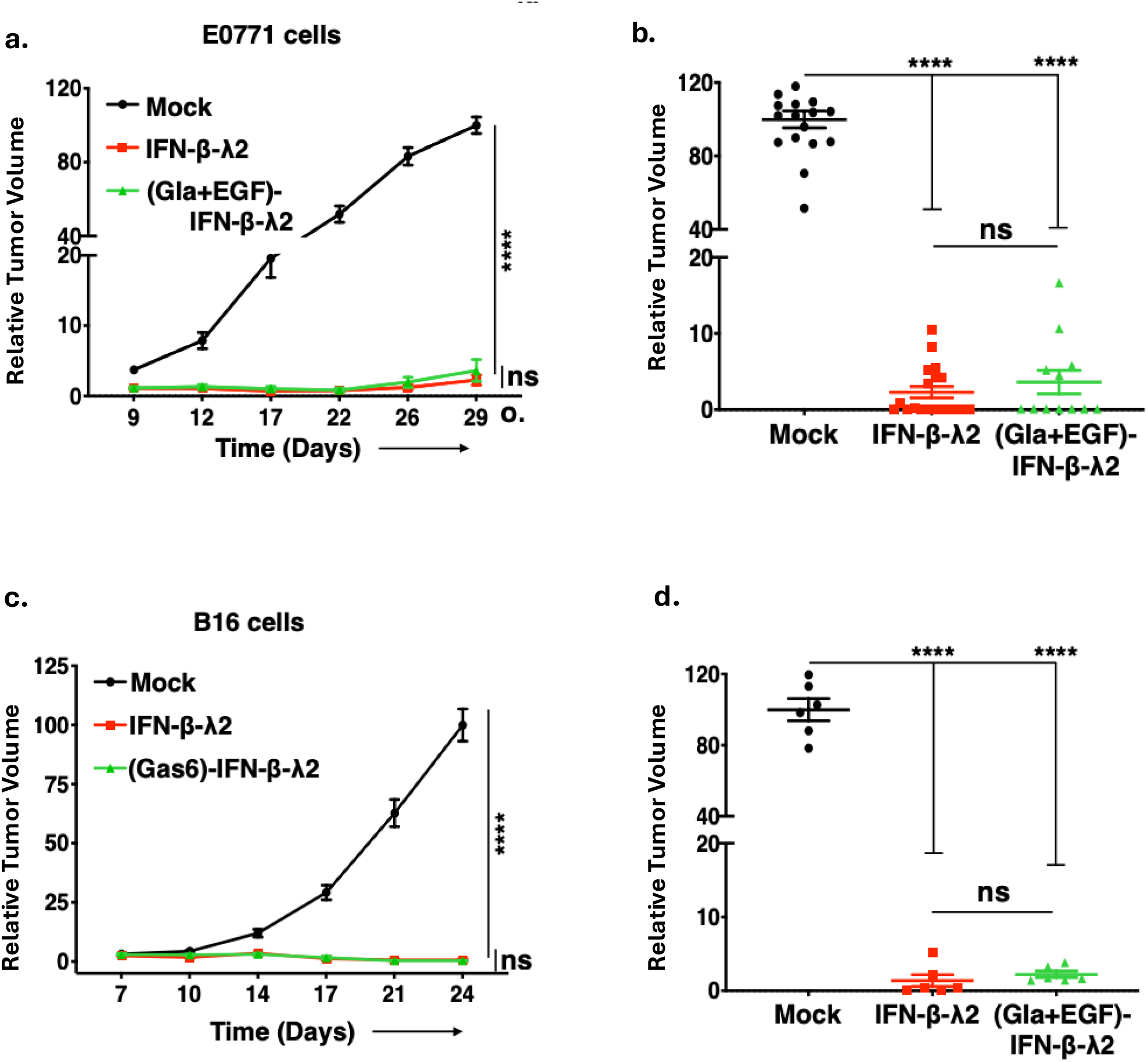
Addition of the Gla and EGF domains do not interfere with anti-tumor activity. To compare the effect of PS targeting on anti-tumor potency of IFN-β-IFN-λ fusion molecule, Gas6(Gla+EGF)-IFN-β-IFN-λ and IFN-β-IFN-λ were compared head to head in E0771 **(a, b)** and B16-F10 models **(c, d).** 5 X 10^4^ E0771 or 5 X 10^4^ B16 cells, mock-transfected or constitutively secreting either IFN-b-IFN-λ or Gas6(Gla+EGF)-IFN-β-IFN-λ, were injected either orthotopically into the mammary fat pad of female C57BL/6 mice for the E0771 model or into the right flank of C57BL/6 mice for the B16-F10 model.

### IFN-β-IFN-λ and Gas6-IFN-β-IFN-λ fusion molecules modulate host immune response to exert anti-tumor activity

To investigate the extrinsic role of tumor microenvironment in the above-mentioned anti-tumor responses, we utilized type I, type III or type I and type III double IFN receptor knockout (KO) mice on C57BL/6 background in E0771 and B16-F10 tumor models (**Fig. 7.a**). As shown in **Fig. 7.b.** (E0771) and **Fig. 7.c.** (B16-F10), when cells expressing IFN-β-IFN-λ or Gas6-IFN-β-IFN-λ were transplanted in IFNAR1 KO mice, tumor growth was partially reduced compared to mock-transfected cells. However, compared to tumor growth in the WT C57BL/6 mice, tumors grew substantially better in the IFNAR1 KO suggesting that type I IFN receptors on tumor-infiltrating immune cells provide a degree of tumor immune surveillance and anti-tumor immunity. In contrast, when IFN-β-IFN-λ or Gas6-IFN-β-IFN-λ expressing tumor cells were transplanted into IFNLR1 KO mice, there was little observable anti-tumor protection (**Fig. 7.d, 7.e**). Similarly, when IFN-β-IFN-λ or Gas6-IFN-β-IFN-λ expressing cells were transplanted into IFNAR1/IFNLR1 double KO mice, the effects of IFNs on tumor growth kinetics were similar to the effects observed in the IFNAR1 single KO mice (**Fig. 7.f, 7.g**). This suggests that type I and type III IFNs act distinctly (likely on different cell types in the TME) and that IFN-β-IFN-λ fusion proteins, as predicted in their design, act on both immune cells (type I IFNs) and on the tumor cells (type III IFNs, see also **Fig. 7.c.**). Consistent with this idea, when IFN-β-IFN-λ or Gas6-IFN-β-IFN-λ expressing tumor cells were transplanted to NOD/SCID mice (E0771 were used in these experiments), tumor growth was only partially suppressed (**Fig.7.h.)** However, when the when IFN-β-IFN-λ or Gas6-IFN-β-IFN-λ expressing E0771 cells were injected into Rag1 KO mice, which are deficient in mature T and B cell cells, all the tumors grew at a similar rate (**Fig.7.i**), suggesting that these fusion proteins exert their anti-tumor effects through the adaptive immune system. Collectively, these data suggest that type I and type III fusion proteins are likely to act to both increase the immunogenicity of tumor cells as well as stimulate immune cells in the tumor microenvironment.

**Figure 7:**
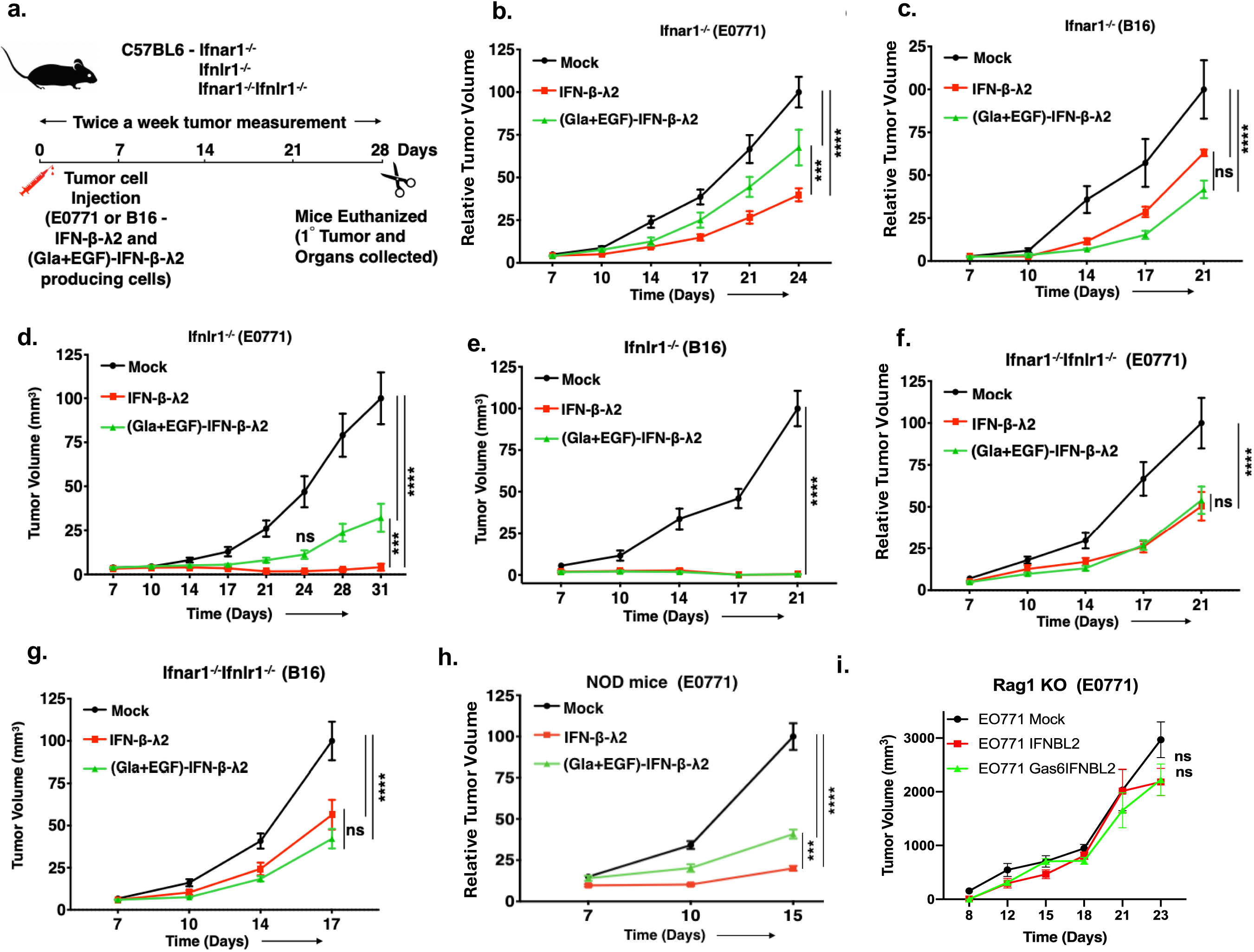
Novel fusion proteins exert anti-tumor responses through the host immune system. **(a)** The graphical outline showing experimental outline of the E0771 and B16-F10 tumor model in type I, type III or type I and type III double IFN receptor KO mice on C57BL/6 background is schematically illustrated. **b-c.** E0771**(b)** and B16-F10 **(c)** mock or constitutively IFN-β-IFN-λ or Gas6-IFN-β-IFN-λ secreting tumor cells were injected in to type I IFN receptor (IFNAR1) KO mice and tumor growth was analyzed. **d-e.** Anti-tumor effects of IFN-β-IFN-λ or Gas6-IFN-β-IFN-λ fusion molecules were also evaluated in type III IFN receptor (IFNLR1) KO mice by monitoring tumor growth dynamics in KO mice injected with mock or IFN fusion molecule secreting E0771 **(d)** and B16-F10 **(e)** tumor cells. **f-g.** Partial inhibition of tumor growth was also observed when E0771 **(f)** and B16-F10 **(g)** mock or constitutively IFN-β-IFN-λ or Gas6-IFN-β-IFN-λ secreting tumor cells were injected into type I and type III IFN receptor double KO mice **(h).** Mock or constitutively IFN-β-IFN-λ or Gas6-IFN-β-IFN-λ secreting E0771 cells were injected into NOD mice and tumor growth analyzed. **(i)** Mock or constitutively IFN-β-IFN-λ or Gas6-IFN-β-IFN-λ secreting E0771 cells were injected into RAG1 KO mice and tumor growth analyzed.

### Novel fusion molecules have direct anti-tumor effects on tumor cells

To explore cell intrinsic versus cell extrinsic functions of IFN-β-IFN-λ and Gas6(Gla+EGF)-IFN-β-IFN-λ (subsequently designated as Gas6-IFN-β-IFN-λ) fusion proteins in the E0771 and B16-F10 models of tumor growth, we assessed the induction of cell surface MHC class I protein expression. Bulk RNA sequencing of primary murine lung epithelial (MLE) cells treated with Gas6(Gla+EGF)-IFN-β-IFN-λ VitK for 7 or 24 hours showed differential regulation of early and late induced genes genes compared to untreated **(Fig. 8.a.)**. Quality control was done by PCA which showed that replicates cluster together and the Gas6(Gla+EGF)-IFN-β-IFN-λ VitK treated samples differ from untreated samples significantly **(Supplementary Fig. 1.e.).** Focusing on genes upregulated at the early time point of 7 hours after treatment, MHC I protein complex and transporter associated antigen processing proteins, Tap1 and Tap2 (**Fig.8.b**.) are seen to be upregulated. Classical ISGs (in green) and PD-L1 (CD274 in red) are also shown to be upregulated (**Fig.8.b**.). Gene ontology enrichment analysis for cellular components identified MHC I protein complexes as most significantly upregulated **(Fig.8.c.)**, whereas regulation of cytokine responses, and pathogen response pathways were the most enriched biological processes **(Fig.8.d.)**. While neither protein had notable effects in cell growth or apoptosis (data not shown), both proteins strongly up-regulated levels of MHC class I protein and PD-L1 expression on the surface of E0771 cells (**Fig. 8.e.,g.**) and B16-F10 cells (**Fig. 8.f.,h,**), predicting enhanced immunogenicity of the modified tumor cells *in vivo*. Consistent with this idea and to analyze STAT1 phosphorylation as a signature IFN response, we extracted steady state lysates, and as shown in (**Supplementary Fig. 1.f.**) in both E0771 and B16-F10 cells constitutively producing IFN-β-IFN-λ or Gas6-IFN-β-IFN-λ fusion molecules, STAT1 phosphorylation was robustly stimulated as compared to mock cells. Taken together, these data support the proof of concept for the utility of PS-targeting IFNs, potentially in combination with checkpoint inhibitors such as anti-PD1.

**Figure 8.**
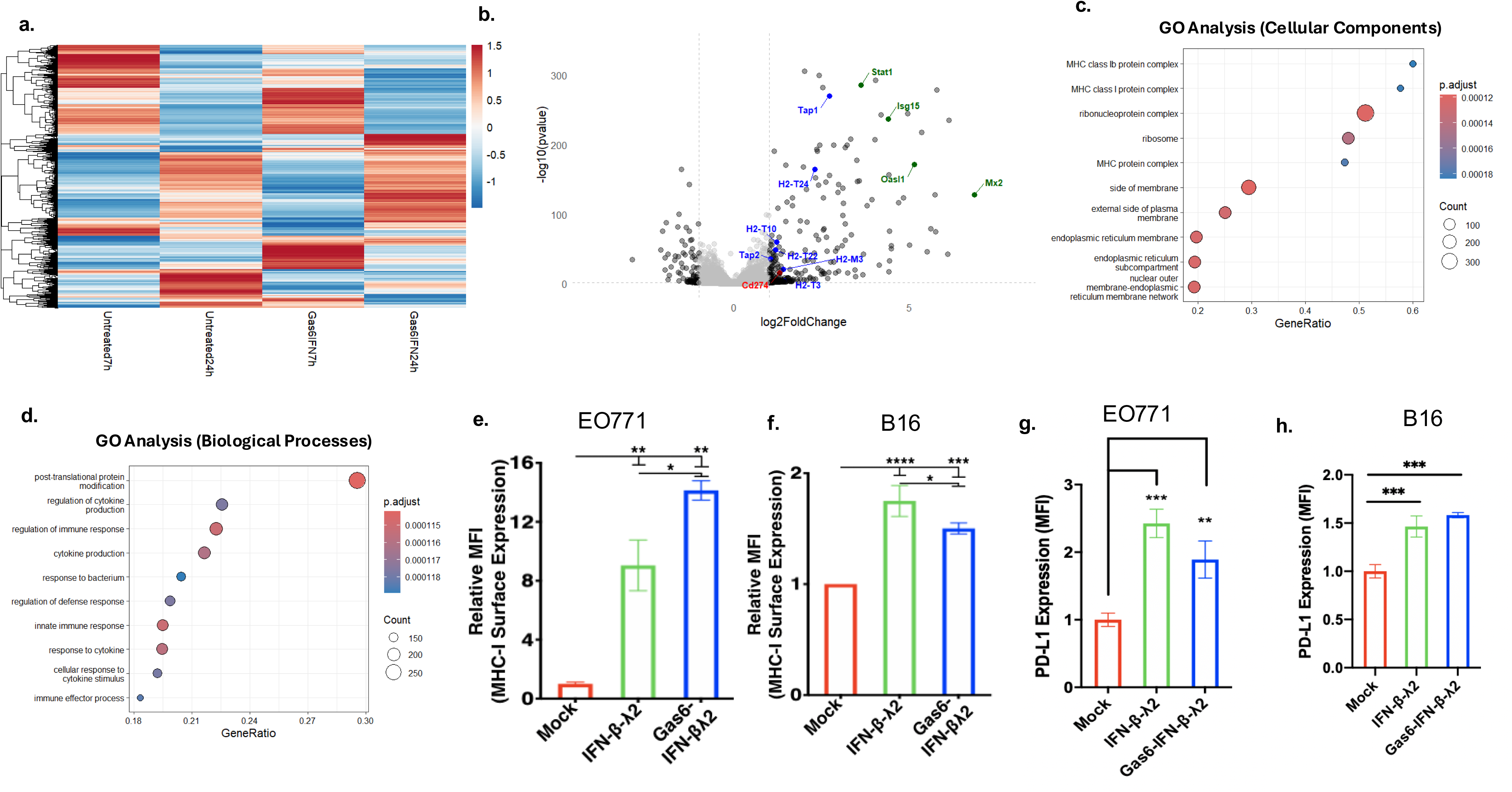
Novel fusion molecules have direct anti-tumor effects on tumor cells. **(a)** RNA sequencing data shows differentially expressed genes upon treatment of MLE cells with Gas6-IFN-β-IFN-λ (VitK) for 7 or 24 hours. **(b)** MHC I, Tap proteins, PD-L1 and classical ISGs are upregulated upon IFN treatment. **(c)** Gene Ontology enrichment revealed that MHC class I protein complexes are upregulated in cells treated with Gas6-IFN-β-IFN-λ (VitK) for 7 hours and **(d)** biological processes related to pathogen response and cytokine production are also upregulated. E0771 and B16-F10 cells mock-transfected or constitutively secreting either IFN-β-IFN-λ or Gas6-IFN-β-IFN-λ were analyzed for cell surface expression of MHC class I (H-2Kd/H-2Dd) protein and PD-L1 by flow cytometry. The data shown are representative of 3 independent experiments (**e-h).** The quantification of relative MFI is shown in MHC-I and PD-L1 respectively (**e,g**) for E0771 and (**f,h**) for B16-F10 cells.

## Discussion

The importance of membrane asymmetry and the redistribution of PS to the outer leaflet of the plasma membrane has gained much interest in the field of immune-oncology in recent years. Unlike PS externalization on stochastic apoptotic cells that are efficiently cleared by macrophage efferocytosis to maintain homeostasis and tolerance, apoptosis in tumors is chronic and sustained, and many solid tumors exist with high apoptotic indexes (tumors have been called wounds that never heal)^23^ ^12^. Similarly, unlike the rapid and reversible externalization of PS on activated immune cells and in the tumor stroma, PS externalization on viable cells in the tumor microenvironment is also chronic and sustained, leading to the robust activation of inhibitory PS receptors in the tumor microenvironment, as well as exhaustion and anergy of immune cells^59^. Such complex sustained PS externalization in the tumor microenvironments is a pathophysiological mechanism that contributes to tumor immune escape by recent studies showing that activation of PS flippases such as TMEM30A can recapitulate immune dysfunction and evasion of host immunity, while blockage of PS externalization improves host immunity^60^. While previous attempts to block PS in the tumor microenvironment using PS-targeting mAbs and PS binding modalities have shown promising results, few if any of these approaches has linked an immunogenic payload to a PS-targeting strategy^61^. Here, we report on a new class of PS targeting modality that fuses a type I and type III IFN duet to the phosphatidylserine binding domain of Gas6.

Type I and type III IFNs bind to different receptor complexes, namely type I (IFNAR) and type III (IFNLR) IFN receptors respectively^62^. While both type I and type III IFN receptors activate a common post-receptor Jak-STAT signaling pathway, recent studies reveal that receptor expression patterns are functional compartmentalized^58^ ^63^. Here, we demonstrate that recombinant fusion proteins that combine type I (IFN-β) and type III (IFN-λ) enhance both antiviral and anti-tumor activity compared to monomeric IFN species, and that this activity is retained when IFN-β-IFN-λ is further linked to the PS-binding domain of Gas6. As such, our approaches highlight the functional co-operativity of type I and type III IFNs as therapeutic entities, and that this synergy can be harnessed using single chain chimeric fusion proteins. While previous studies have shown that co-administration of IFN-α and IFN-λ provides synergistic antitumor effect in the mouse model of hepatocellular carcinoma ^55^, likely attributed to distinct patterns of cell type and tissue distribution of the IFN receptors, the present approach permits focal localization of IFNs and potentially reduced side effects of type I IFNs. Moreover, the findings that IFN-β-IFN-λ retains full activity when fused to a PS targeting Gla domain posits that such molecules can be used to specifically target areas of high local PS exposure, such as the microenvironments of solid tumors as well as in areas of viral infection. The present study provides a proof-of-concept milestone for the use of functional gain-of-function chimeric type I and type III IFN fusion proteins for potential therapeutic applications.

The idea that type I and type III IFNs will target both immune cells and epithelial to improve efficacy is predicted by the differences in tissue distribution of IFNAR and IFNLR. The concept that fusing IFN-β-IFN-λ to the Gla and EGF-like domains of Gas6 is expected to add a level of localized action to IFNs to target specific areas of stressed and diseased cells and exploit a fortuitous vulnerability in cancer and virology as noted above ^12^ ^24^ ^64^. Two of the better understood strategies to target PS include recombinant Annexin V proteins ^65^ ^26^ ^66^ and PS targeting mAbs generated in the Thorpe laboratory and subsequently developed into preclinical and clinical utilities by Peregerine Pharmaceuticals ^67^ ^68^ ^27^ ^69^. In the case for recombinant Annexin V, previous studies have shown that Annexin V, when administered *in vivo*, can improve host anti-tumor immunity, presumably by blocking immunosuppressive signals from PS ^13^. Annexin V also renders apoptotic tumor cells immunogenic, the addition of Annexin V to apoptotic tumor cell vaccines increased the percentage of tumor-free mice in syngeneic tumor cure assays ^70^. More recently, administration of Annexin V was shown to significantly enhance immunogenicity and anti-tumor efficacy when administered with chemotherapy and further showed synergy with immune checkpoint inhibitors^66^. Analogous to the biology of Annexin V, PS targeting mAbs, including Bavituximab and mch1N11 have also showed promising pre-clinical activities, often leading to increased immunogenicity in tumors, increased influx of CD4 and CD8 T cells, and synergy with immune checkpoint inhibitors and radiation therapy^27^ ^14^. In recent years, the repertoire of PS targeting approaches has gained further traction including the development of PS targeting agents as cationic small molecules (QW4869)^71^, β2GPI binding F(ab)2 fusion proteins^72^, Annexin fusion proteins (to cystathione gamma-lyase)^73^, nanovesicles that target PS (SapC-DOPS)^31^ ^74^, and peptide-peptoid hybrids (PPS1D1) ^75^. While such strategies have versability to modify payloads, most of the current PS-targeting strategies aim to target PS in stressed/diseased tissues, they lack an immunogenic payload. The Gas6-IFN fusions described here not on directly target PS but divert immunosuppressive signals into immunogenic signals to enhance antiviral and anti-tumor immune responses.

While the present study shows feasibility and proof-of-concept assurance that trimeric fusions, when produced locally in the tumor microenvironment, retain biological activates as active IFNs with PS-targeting capacity, future studies will be required to optimize their *in vivo* delivery and whether they can be injected systemically, or intra-tumorally. For PS-targeting mAbs, such as Bavituximab, systemic infusion of mAbs lead to honing and localization in the tumor microenvironment. We also expect that Gas6-IFNs can be applied systemically, since Gas6 has nanomolar affinity toward PS, and Gla domains of Gas6 have been shown to cluster and/or oligomerize around PS that is constitutively externalized in the tumor microenvironment. In sum, we hypothesize that the targeting of Gas6-IFNβ-IFNλ will have an important role in activating host anti-tumor immunity. Indeed, this strategy is in line with emerging concepts that PS in the tumor microenvironment is immunosuppressive and that type I and type III IFNs have dual compartmentalized models of immune activation. Moreover, since Gas6-IFNβ-IFNλ induces MHC class I and PD-L1 on the PS-positive cells, these molecules might prove beneficial via the combined effects of checkpoint therapeutics such as anti-PD1 or anti-PD-L1.

## Acknowledgements

We thank members of our laboratory for helpful discussions. This work was supported in part by NIH R01 CA260137-01A1 from the National Cancer Institute to RBB and SVK. Biorender was used to create figures. RBB and SVK disclose they are cofounders of a biotechnology company called Targeron Therapeutics that aims to develop PS-targeting IFNs for anti-cancer and anti-viral applications.

## Supplementary figures

**Supplementary figure 1.**
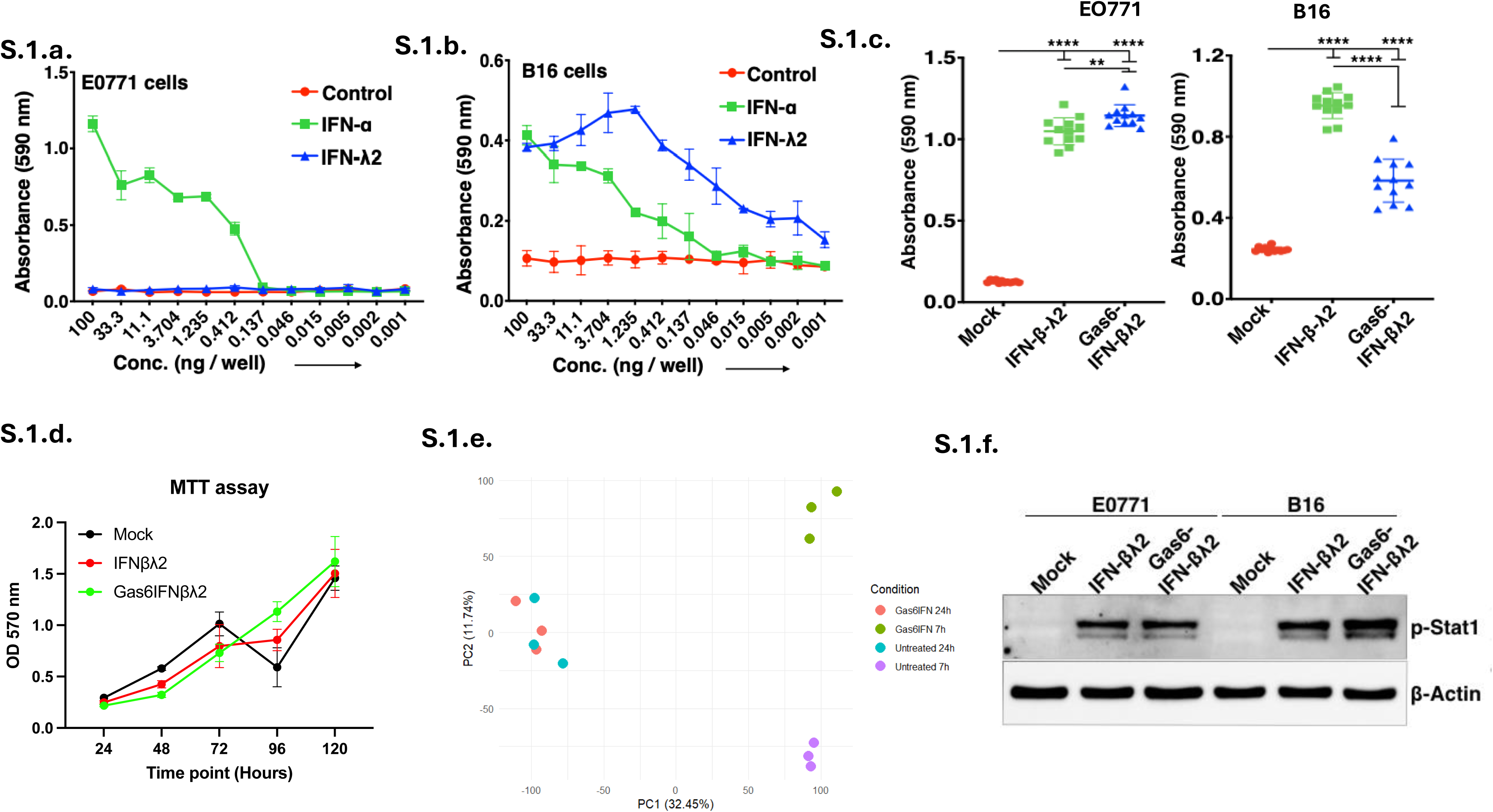
Antiviral assay with **(a)** E0771 and **(b)** B16 cells shows that E0771 cells respond only to type I IFNs (IFN-α) and B16 cells respond to both type I IFNs (IFN-α) and type III IFNs (IFN-λ). **(c)** Shows protection anti-viral from VSV-GFP using IFN fusion proteins in EO771 and B16 cells, as shown by crystal violet staining. **(d)** MTT assay shows no significant differences in cell viablility of E0771 cells producing IFN fusion proteins **(e)** Principal component analysis of RNA sequencing data from cells treated with Gas6(Gla+EGF)-IFN-λ2(VitK) for 7 or 24 hours shows clustering of samples. **(f)** IFN fusion proteins activate IFN signaling, as seen by upregulation of p-STAT-1 in E0771 and B16 cells.

**Supplementary figure 2.**
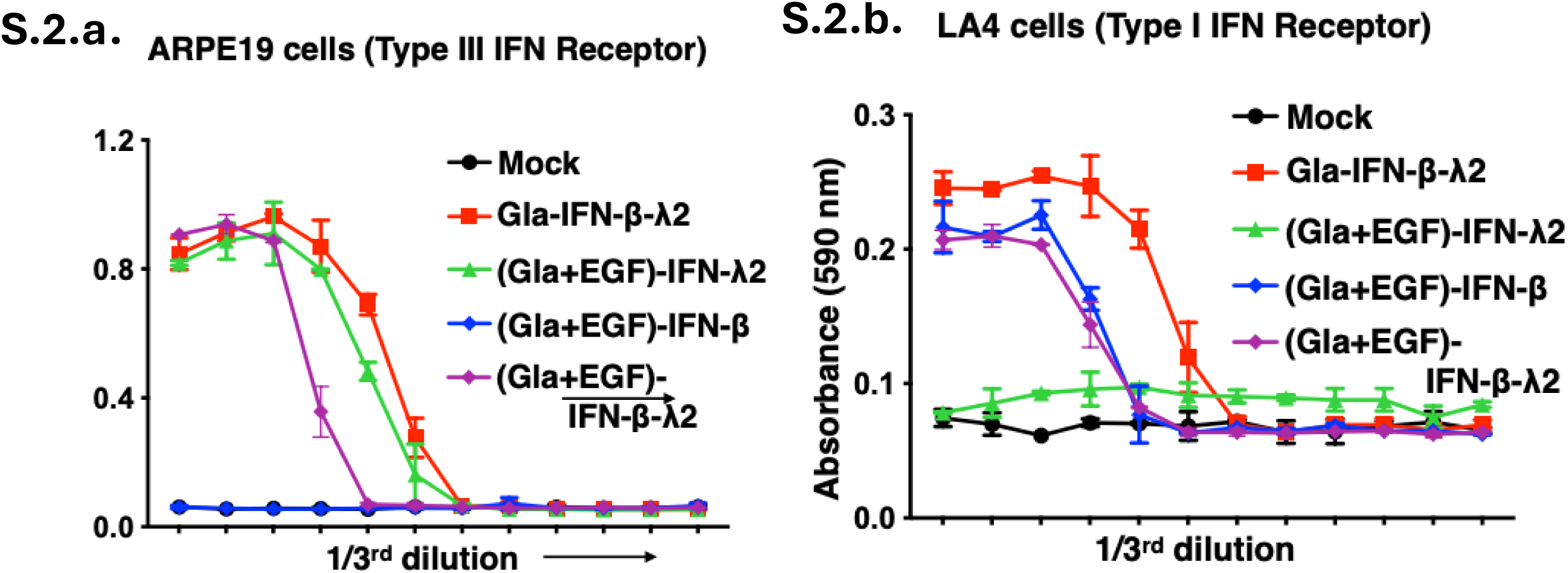
Antiviral activities of Gas6(Gla)-IFN-β-IFN-λ, Gas6(Gla+EGF)-IFN-λ2, Gas6(Gla+EGF)-IFN-β or Gas6(Gla+EGF)-IFN-β-IFN-λ protein containing HEK293T supernatants are measured in **(a)** LA4 and **(b)** ARPE19 cells.

**Supplementary figure 3.**
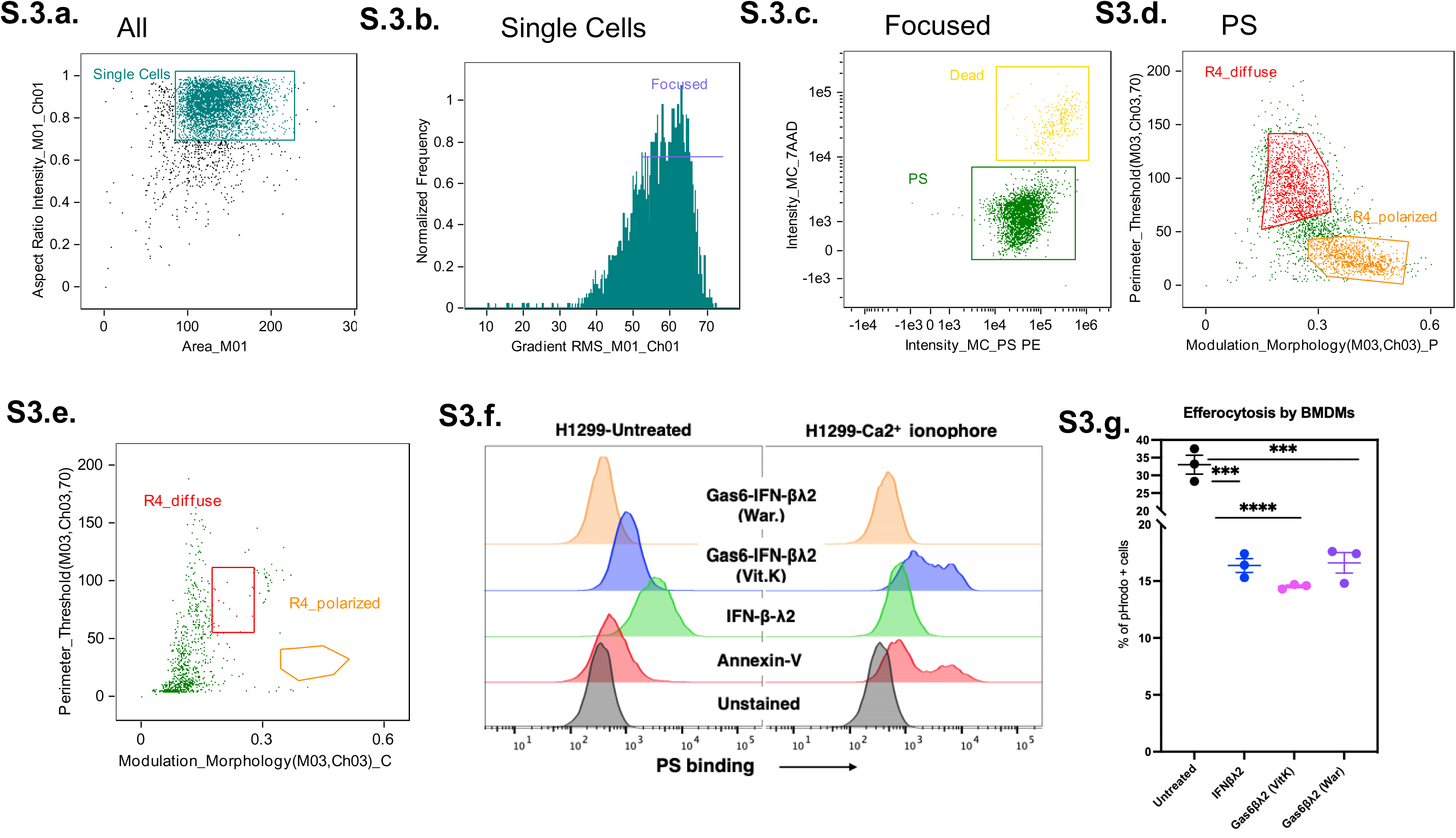
Gating scheme for the imaging flow cytometry done in Figure 4. Single cells were gated by area and aspect ratio of the events collected **(a,b).** Live cells were gated (green) by excluding dead cells (yellow) that stained positive for 7AAD **(c)**. The surface staining pattern was quantified by measuring the perimeter staining and the morphology **(d)**. Gas6(Gla+EGF)-IFN-β-IFN-λ (War) did not show binding to CDC50AED29 cells **(e). (f)** H1299 cancer cells were treated with A23187 to induce calcium stress mediated PS exposure. Only Gas6(Gla+EGF)-IFN-β-IFN-λ (VitK) binds to PS^+^ cells, like Annexin V that was used as a positive control for binding to externalize PS. IFN fusion proteins blocked efferocytosis of apoptotic cells by bone marrow derived macrophages **(g)**.

**Supplementary Table 1.**
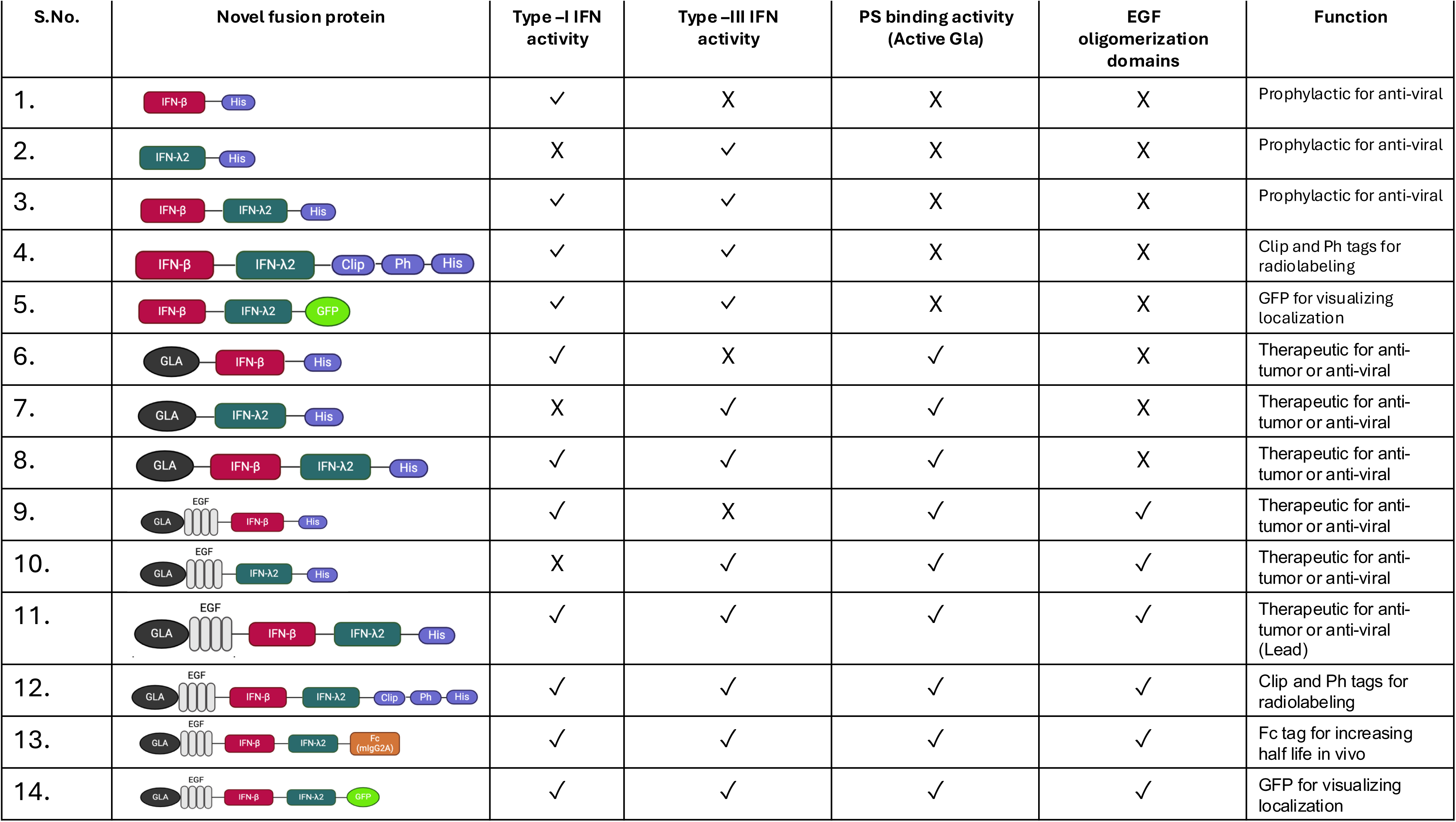
Shows the iterations of IFN fusion proteins that can be used as for developing pre-clinical models and for therapeutics.

| S.No. | Novel fusion protein | Type –I IFN activity | Type –III IFN activity | PS binding activity (Active Gla) | EGF oligomerization domains | Function |
| --- | --- | --- | --- | --- | --- | --- |
| 1. |  | ✓ | X | X | X | Prophylactic for anti-viral |
| 2. |  | X | ✓ | X | X | Prophylactic for anti-viral |
| 3. |  | ✓ | ✓ | X | X | Prophylactic for anti-viral |
| 4. |  | ✓ | ✓ | X | X | Clip and Ph tags for radiolabeling |
| 5. |  | ✓ | ✓ | X | X | GFP for visualizing localization |
| 6. |  | ✓ | X | ✓ | X | Therapeutic for anti-tumor or anti-viral |
| 7. |  | X | ✓ | ✓ | X | Therapeutic for anti-tumor or anti-viral |
| 8. |  | ✓ | ✓ | ✓ | X | Therapeutic for anti-tumor or anti-viral |
| 9. |  | ✓ | X | ✓ | ✓ | Therapeutic for anti-tumor or anti-viral |
| 10. |  | X | ✓ | ✓ | ✓ | Therapeutic for anti-tumor or anti-viral |
| 11. |  | ✓ | ✓ | ✓ | ✓ | Therapeutic for anti-tumor or anti-viral (Lead) |
| 12. |  | ✓ | ✓ | ✓ | ✓ | Clip and Ph tags for radiolabeling |
| 13. |  | ✓ | ✓ | ✓ | ✓ | Fc tag for increasing half life in vivo |
| 14. |  | ✓ | ✓ | ✓ | ✓ | GFP for visualizing localization |

## References

1. Agarwal, D., et al., Advances in Vaccines, Checkpoint Blockade, and Chimeric Antigen Receptor-Based Cancer Immunotherapeutics. Crit Rev Immunol, 2025. 45(1): p. 65–80.

2. Sharma, P., et al., Immune checkpoint therapy-current perspectives and future directions. Cell, 2023. 186(8): p. 1652–1669.

3. Sharma, P., et al., The Next Decade of Immune Checkpoint Therapy. Cancer Discov, 2021. 11(4): p. 838–857.

4. Budczies, J., et al., Tumour mutational burden: clinical utility, challenges and emerging improvements. Nat Rev Clin Oncol, 2024. 21(10): p. 725–742.

5. Fares, C.M., et al., Mechanisms of Resistance to Immune Checkpoint Blockade: Why Does Checkpoint Inhibitor Immunotherapy Not Work for All Patients? Am Soc Clin Oncol Educ Book, 2019. 39: p. 147–164.

6. Labani-Motlagh, Alireza, Mehrnoush Ashja-Mahdavi, and Angelica Loskog. ‘The tumor microenvironment: a milieu hindering and obstructing antitumor immune responses.’ Frontiers in immunology 11 (2020): 940.

7. Kroemer, G., et al., Immunogenic cell stress and death. Nat Immunol, 2022. 23(4): p. 487–500.

8. Kroemer, G., Galluzzi, L., Kepp, O. & Zitvogel, L. Immunogenic Cell Death in Cancer Therapy. Annu. Rev. Immunol. 31, 51–72 (2013).

9. Wu, B., et al., Cold and hot tumors: from molecular mechanisms to targeted therapy. Signal Transduct Target Ther, 2024. 9(1): p. 274.

10. Ibis, B., et al., Immune-related adverse effects of checkpoint immunotherapy and implications for the treatment of patients with cancer and autoimmune diseases. Front Immunol, 2023. 14: p. 1197364.

11. Leventis, P.A. and S. Grinstein, The distribution and function of phosphatidylserine in cellular membranes. Annu Rev Biophys, 2010. 39: p. 407–27.

12. Birge, R. B. et al. Phosphatidylserine is a global immunosuppressive signal in efferocytosis, infectious disease, and cancer. Cell Death Differ. 23, 962–978 (2016).

13. Bondanza, A., et al., Inhibition of phosphatidylserine recognition heightens the immunogenicity of irradiated lymphoma cells in vivo. J Exp Med, 2004. 200(9): p. 1157–65.

14. Budhu, S. et al. Targeting Phosphatidylserine Enhances the Anti-tumor Response to Tumor-Directed Radiation Therapy in a Preclinical Model of Melanoma. Cell Rep. 34, 108620 (2021).

15. Ran, S., Downes, A. & Thorpe, P. E. Increased Exposure of Anionic Phospholipids on the Surface of Tumor Blood Vessels. Cancer Res. 62, 6132–6140 (2002).

16. Ran, S. and P.E. Thorpe, Phosphatidylserine is a marker of tumor vasculature and a potential target for cancer imaging and therapy. Int J Radiat Oncol Biol Phys, 2002. 54(5): p. 1479–84.

17. Sakuragi, Takaharu, and Shigekazu Nagata. ‘Regulation of phospholipid distribution in the lipid bilayer by flippases and scramblases.’ Nature Reviews Molecular Cell Biology 24.8 (2023): 576–596.

18. Nagata, S., Sakuragi, T. & Segawa, K. Flippase and scramblase for phosphatidylserine exposure. Curr. Opin. Immunol. 62, 31–38 (2020).

19. Suzuki, J., et al., Xk-related protein 8 and CED-8 promote phosphatidylserine exposure in apoptotic cells. Science, 2013. 341(6144): p. 403–6.

20. Suzuki, J., Imanishi, E. & Nagata, S. Xkr8 phospholipid scrambling complex in apoptotic phosphatidylserine exposure. Proc. Natl. Acad. Sci. 113, 9509–9514 (2016).

21. Ryoden, Y., K. Segawa, and S. Nagata, Requirement of Xk and Vps13a for the P2X7-mediated phospholipid scrambling and cell lysis in mouse T cells. Proc Natl Acad Sci U S A, 2022. 119(7).

22. Suzuki, J., Umeda, M., Sims, P. J. & Nagata, S. Calcium-dependent phospholipid scrambling by TMEM16F. Nature 468, 834–838 (2010).

23. Gadiyar, V., et al., Cell Death in the Tumor Microenvironment: Implications for Cancer Immunotherapy. Cells, 2020. 9(10).

24. Kumar, S., D. Calianese, and R.B. Birge, Efferocytosis of dying cells differentially modulate immunological outcomes in tumor microenvironment. Immunol Rev, 2017. 280(1): p. 149–164.

25. Behuria, Himadri Gourav, Sabyasachi Dash, and Santosh Kumar Sahu. ‘Phospholipid scramblases: Role in cancer progression and anticancer therapeutics.’ Frontiers in Genetics 13 (2022): 875894.

26. Stach, C.M., et al., Treatment with annexin V increases immunogenicity of apoptotic human T-cells in Balb/c mice. Cell Death Differ, 2000. 7(10): p. 911–5.

27. Gray, M.J., et al., Phosphatidylserine-targeting antibodies augment the anti-tumorigenic activity of anti-PD-1 therapy by enhancing immune activation and downregulating pro-oncogenic factors induced by T-cell checkpoint inhibition in murine triple-negative breast cancers. Breast Cancer Res, 2016. 18(1): p. 50.

28. Desai, Tanvi J., et al. ‘Identification of lipid-phosphatidylserine (PS) as the target of unbiasedly selected cancer specific peptide-peptoid hybrid PPS1.’ Oncotarget 7.21 (2016): 30678.

29. Lee, Gyeongwoo, et al. ‘Development of Apoptotic-Cell-Inspired Antibody–Drug Conjugate for Effective Immune Modulation.’ International Journal of Molecular Sciences 24.22 (2023): 16036.

30. Li, Ran, et al. ‘Targeting Phosphatidylserine with Calcium-Dependent Protein–Drug Conjugates for the Treatment of Cancer.’ Molecular cancer therapeutics 17.1 (2018): 169–182.

31. Davis, H.W., et al., Biotherapy of Brain Tumors with Phosphatidylserine-Targeted Radioiodinated SapC-DOPS Nanovesicles. Cells, 2020. 9(9).

32. Pestka, S., et al., The interferon gamma (IFN-gamma) receptor: a paradigm for the multichain cytokine receptor. Cytokine Growth Factor Rev, 1997. 8(3): p. 189–206.

33. Samuel, C.E., Antiviral actions of interferon. Interferon-regulated cellular proteins and their surprisingly selective antiviral activities. Virology, 1991. 183(1): p. 1–11.

34. Mesev, E.V., R.A. LeDesma, and A. Ploss, Decoding type I and III interferon signalling during viral infection. Nat Microbiol, 2019. 4(6): p. 914–924.

35. Platanias, L.C., Mechanisms of type-I- and type-II-interferon-mediated signalling. Nat Rev Immunol, 2005. 5(5): p. 375–86.

36. Park, A. and A. Iwasaki, Type I and Type III Interferons - Induction, Signaling, Evasion, and Application to Combat COVID-19. Cell Host Microbe, 2020. 27(6): p. 870–878.

37. Stanifer, M.L., K. Pervolaraki, and S. Boulant, Differential Regulation of Type I and Type III Interferon Signaling. Int J Mol Sci, 2019. 20(6).

38. Donnelly, R.P., et al., The expanded family of class II cytokines that share the IL-10 receptor-2 (IL-10R2) chain. J Leukoc Biol, 2004. 76(2): p. 314–21.

39. Kotenko, S.V., et al., IFN-λambdas mediate antiviral protection through a distinct class II cytokine receptor complex. Nat Immunol, 2003. 4(1): p. 69–77.

40. Sheppard, P., et al., IL-28, IL-29 and their class II cytokine receptor IL-28R. Nat Immunol, 2003. 4(1): p. 63–8.

41. Schneider, W.M., M.D. Chevillotte, and C.M. Rice, Interferon-stimulated genes: a complex web of host defenses. Annu Rev Immunol, 2014. 32: p. 513–45.

42. Stark, G.R. and J.E. Darnell, Jr., The JAK-STAT pathway at twenty. Immunity, 2012. 36(4): p. 503–14.

43. Lazear, H.M., T.J. Nice, and M.S. Diamond, Interferon-lambda: Immune Functions at Barrier Surfaces and Beyond. Immunity, 2015. 43(1): p. 15–28.

44. Lazear, H.M., J.W. Schoggins, and M.S. Diamond, Shared and Distinct Functions of Type I and Type III Interferons. Immunity, 2019. 50(4): p. 907–923.

45. Zhou, J.H., et al., Type III Interferons in Viral Infection and Antiviral Immunity. Cell Physiol Biochem, 2018. 51(1): p. 173–185.

46. Sistigu, A., et al., Cancer cell-autonomous contribution of type I interferon signaling to the efficacy of chemotherapy. Nat Med, 2014. 20(11): p. 1301–9.

47. Zitvogel, L., et al., Type I interferons in anticancer immunity. Nat Rev Immunol, 2015. 15(7): p. 405–14.

48. Musella, M., et al., Type-I-interferons in infection and cancer: Unanticipated dynamics with therapeutic implications. Oncoimmunology, 2017. 6(5): p. e1314424.

49. Thyrell, L., et al., Mechanisms of Interferon-alpha induced apoptosis in malignant cells. Oncogene, 2002. 21(8): p. 1251–62.

50. Corrales, L., et al., The host STING pathway at the interface of cancer and immunity. J Clin Invest, 2016. 126(7): p. 2404–11.

51. Garg, A.D. and P. Agostinis, Cell death and immunity in cancer: From danger signals to mimicry of pathogen defense responses. Immunol Rev, 2017. 280(1): p. 126–148.

52. Sprooten, J., P. Agostinis, and A.D. Garg, Type I interferons and dendritic cells in cancer immunotherapy. Int Rev Cell Mol Biol, 2019. 348: p. 217–262.

53. Lasfar, A., et al., Characterization of the mouse IFN-λambda ligand-receptor system: IFN-λambdas exhibit antitumor activity against B16 melanoma. Cancer Res, 2006. 66(8): p. 4468–77.

54. Li, W., et al., Type III interferon induces apoptosis in human lung cancer cells. Oncol Rep, 2012. 28(3): p. 1117–25.

55. Lasfar, A., et al., Concerted action of IFN-alpha and IFN-λambda induces local NK cell immunity and halts cancer growth. Oncotarget, 2016. 7(31): p. 49259–49267.

56. Souza-Fonseca-Guimaraes, Fernando, et al. ‘NK cells require IL-28R for optimal in vivo activity.’ Proceedings of the National Academy of Sciences 112.18 (2015): E2376–E2384.

57. Burkart, Christoph, et al. ‘Usp18 deficient mammary epithelial cells create an antitumour environment driven by hypersensitivity to IFN-λ and elevated secretion of Cxcl10.’ EMBO molecular medicine 5.7 (2013): 1035–1050.

58. McElrath, C., Espinosa, V., Lin, J. D., Peng, J., Sridhar, R., Dutta, O., … & Kotenko, S. V. (2021). Critical role of interferons in gastrointestinal injury repair. Nature communications, 12(1), 2624.

59. Calianese, D. C. & Birge, R. B. Biology of phosphatidylserine (PS): basic physiology and implications in immunology, infectious disease, and cancer. Cell Commun. Signal. 18, 41 (2020).

60. Wang, Weihong, et al. ‘Mobilizing phospholipids on tumor plasma membrane implicates phosphatidylserine externalization blockade for cancer immunotherapy.’ Cell reports 41.5 (2022).

61. Huang X, Ye D, Thorpe PE. Enhancing the potency of a whole-cell breast cancer vaccine in mice with an antibody-IL-2 immunocytokine that targets exposed phosphatidylserine. Vaccine. 2011 Jun 24;29(29-30):4785–93. doi: 10.1016/j.vaccine.2011.04.082. Epub 2011 May 8. PMID: 21557977.

62. Levy, D.E., I.J. Marie, and J.E. Durbin, Induction and function of type I and III interferon in response to viral infection. Curr Opin Virol, 2011. 1(6): p. 476–86.

63. Kotenko, S.V. and J.E. Durbin, Contribution of type III interferons to antiviral immunity: location, location, location. J Biol Chem, 2017. 292(18): p. 7295–7303.

64. Geng, K. et al. Requirement of Gamma-Carboxyglutamic Acid Modification and Phosphatidylserine Binding for the Activation of Tyro3, Axl, and Mertk Receptors by Growth Arrest-Specific 6. Front. Immunol. 8, (2017).

65. Munoz, L.E., et al., The role of annexin A5 in the modulation of the immune response against dying and dead cells. Curr Med Chem, 2007. 14(3): p. 271–7.

66. Kang, T. H. et al. Annexin A5 as an immune checkpoint inhibitor and tumor-homing molecule for cancer treatment. Nat. Commun. 11, 1137 (2020).

67. Beck, A.W., et al., Combination of a monoclonal anti-phosphatidylserine antibody with gemcitabine strongly inhibits the growth and metastasis of orthotopic pancreatic tumors in mice. Int J Cancer, 2006. 118(10): p. 2639–43.

68. DeRose, P., P.E. Thorpe, and D.E. Gerber, Development of bavituximab, a vascular targeting agent with immune-modulating properties, for lung cancer treatment. Immunotherapy, 2011. 3(8): p. 933–44.

69. He, J., et al., Antiphosphatidylserine antibody combined with irradiation damages tumor blood vessels and induces tumor immunity in a rat model of glioblastoma. Clin Cancer Res, 2009. 15(22): p. 6871–80.

70. Frey, B., et al., AnnexinA5 renders dead tumor cells immunogenic--implications for multimodal cancer therapies. J Immunotoxicol, 2009. 6(4): p. 209–16.

71. Vuckovic, S., et al., The cationic small molecule GW4869 is cytotoxic to high phosphatidylserine-expressing myeloma cells. Br J Haematol, 2017. 177(3): p. 423–440.

72. Phinney, Natalie Z., et al. ‘Development of betabodies: The next generation of phosphatidylserine targeting agents.’ Journal of Biological Chemistry 300.9 (2024).

73. Krais, J.J., et al., Antitumor Synergism and Enhanced Survival with a Tumor Vasculature-Targeted Enzyme Prodrug System, Rapamycin, and Cyclophosphamide. Mol Cancer Ther, 2017. 16(9): p. 1855–1865.

74. N’Guessan, K.F., P.H. Patel, and X. Qi, SapC-DOPS - a Phosphatidylserine-targeted Nanovesicle for selective Cancer therapy. Cell Commun Signal, 2020. 18(1): p. 6.

75. Matharage, J.M., et al., Unbiased Selection of Peptide-Peptoid Hybrids Specific for Lung Cancer Compared to Normal Lung Epithelial Cells. ACS Chem Biol, 2015. 10(12): p. 2891–9.

